# Instrumental-Motor Transfer: The Relative Value of Competing Movement Goals Modulates Implicit Motor Learning

**DOI:** 10.64898/2026.06.22.733849

**Authors:** Naser Al-Fawakhiri, Vikram Chib, Samuel D. McDougle

## Abstract

Motor learning depends on the integration of several, often competing signals, including reward and punishment (which guide learning the value of actions via reinforcement learning) and sensorimotor errors (which refine the execution of actions via motor adaptation). If and how these signals interact remains controversial. Here we show that the learned value of actions can fundamentally alter how the motor adaptation system learns from errors. Human participants learned the reward values of different movement targets and subsequently underwent implicit motor learning at those targets. Across several experiments, motor learning was robustly attenuated when it brought the limb toward previously punished movement directions. Moreover, learning was globally suppressed when punished targets were themselves the movement goal. Learning was not reciprocally enhanced when it pushed movements toward previously rewarded directions, revealing a valence asymmetry. Using computational modeling, we present a novel mechanism to explain our effects: the motor adaptation system tunes its error sensitivity according to a movement-specific value gradient. Our findings point to a close interaction between reinforcement learning and motor memory — a kind of “instrumental-motor transfer” – where learned action values modulate error-based motor adaptation.

**Significance:** In order to ensure our motor behavior yields our desired outcomes, we must not only learn which actions lead to which outcomes (via reinforcement learning) but also how to precisely execute those actions to achieve the desired outcome (by calibrating our actions via implicit motor adaptation). While it is clear that reinforcement learning and motor adaptation operate at different levels of an action selection hierarchy, it is unclear if they interact. Knowing if and how these processes interact is critical for understanding the fundamental algorithms and various neural circuits underlying real-world motor behavior. Here, we use behavioral data and computational modeling to reveal a novel interaction between the two processes, where motor adaptation is suppressed when it would bring the hand toward a previously punished movement goal.

## Introduction

We move about the world in a complex landscape of movement goals, each with a different learned or intrinsic value. For example, when wandering around the kitchen, moving your hand towards a tasty snack is a much more valuable goal than moving it onto a hot stovetop. This “affordance” landscape for action^1^ is in part shaped by reinforcement learning^2^, where we learn the value of different actions via the rewards or punishments those actions tend to produce, and adjust action selection accordingly^3^.

At a lower level, our motor system must learn how to physically accomplish our movement goals by generating motor commands that produce desired sensory consequences. Performing movements accurately requires unconsciously calibrating motor commands to reduce errors, a process known as motor adaptation^4,5^. While it is clear that these two forms of learning — reinforcement learning and motor adaptation — are distinct and operate at different levels of an action selection hierarchy, it is unclear if they interact.

Some existing evidence in the domain of visuomotor reach adaptation points to an interaction between value-based learning signals (rewards and punishments) and motor adaptation. Galea et al^6^ found that when people learned to counteract visuomotor perturbations to hit a target, the addition of reward and punishment feedback significantly modulated motor learning. Specifically, punishment accelerated early learning while reward seemingly enhanced retention of the learned rotation. This study and several more recent experiments^7–9^ are taken as evidence that the implicit motor adaptation system is sensitive not only to sensory prediction errors (e.g., an unexpected sensory outcome of a movement to a target) but also to motivational or task success signals (e.g., earning $1 when hitting the target). In general, these studies have found that reward or successful feedback diminishes error sensitivity, slowing learning, while punishment or task failure enhances error sensitivity, spurring faster adaptation.

However, other results have complicated the interpretations of these findings. First, recent work has suggested that implicit motor adaptation may not be sensitive to reward feedback^10,11^; instead, reward may act on other learning processes that operate in parallel to motor adaptation, such as deliberate (explicit) aiming strategies ^5,12–17^, which were not fully eliminated in prior studies that reported an effect of reward. Moreover, purported effects of task success or failure on adaptation are susceptible to subtle methodological changes and may be better explained through alternative processes^18^. It is thus still unclear whether value-based learning modulates implicit motor adaptation, or if it affects motor output in other ways, perhaps by reinforcing deliberate strategies or explicit motor memories, or by otherwise biasing the selection of rewarded or successful actions^14,15,19–24^.

Here we took a different approach to this topic: Instead of grafting reward or punishment feedback onto a motor adaptation task, we had people independently learn the value of competing movement goals prior to adaptation. We propose that the relative value of actions creates a value “gradient” that subsequently tunes the learning rate of implicit adaptation for all similar actions. Namely, adaptation around low-valued actions exhibits diminished error sensitivity, slowing learning to avoid producing punished actions. To test this framework, we developed a computational model of adaptation and fit this model to three separate behavioral experiments. In all experiments, participants first completed a movement-based reinforcement learning task^25,26^ followed by a motor learning task that isolates implicit adaptation^27^. We then measured the effect of the relative value of actions on adaptation. Our results validated our computational model, revealing a suppression of adaptation for low-value actions. Such results point to a computational link between the value landscape of our immediate environment and the motor system’s automatic processes of sensorimotor learning.

## Results

### Adaptation is suppressed in the vicinity of punished reach goals

In Experiment 1, we tested the hypothesis that implicit motor adaptation would be suppressed when it brought the effector toward low-valued actions. Participants reached to one of two visual goals (targets) while receiving “clamped” visual error feedback, where the cursor followed a fixed trajectory slightly off center from the goal. This clamped feedback is designed to elicit implicit visuomotor adaptation by producing a discrepancy between the observed cursor trajectory and the intended movement, known as a “sensory-prediction error” (**Fig. 1A**). In the Control Phases (**Fig. 1B**), both targets were neutral, where reaches to both did not result in the gain or loss of any points. This was followed by a Reward Association Phase, which established high-value (i.e., hitting the target led to a gain of points on average) versus low-value (hitting the target led to a loss of points on average) targets. Next, in the Test Phases, participants reached to the higher-value target while receiving clamped feedback. The high-value target was either placed in the location where they had most recently (i.e., in the Reward Association Phase) received rewards (Test Phase 1) or punishment (Test Phase 2). Across consecutive mini-blocks of adaptation, in both Control and Test Phases, the sign of the clamp alternated. This allowed us to measure the degree of adaptation when learning moved the hand *toward* a low-value target (TOWARD condition) versus *away* from a low-value target (AWAY condition). Our design also allowed us to measure if any differences between AWAY and TOWARD conditions were related to specific movement directions, or to more abstract variables (i.e., target identities); that is, participants always reached to the high-value target, but, in Test Phase 2, the location of the high-value target moved to the location where they had recently received punishment in the reward phase.

**Figure 1.**
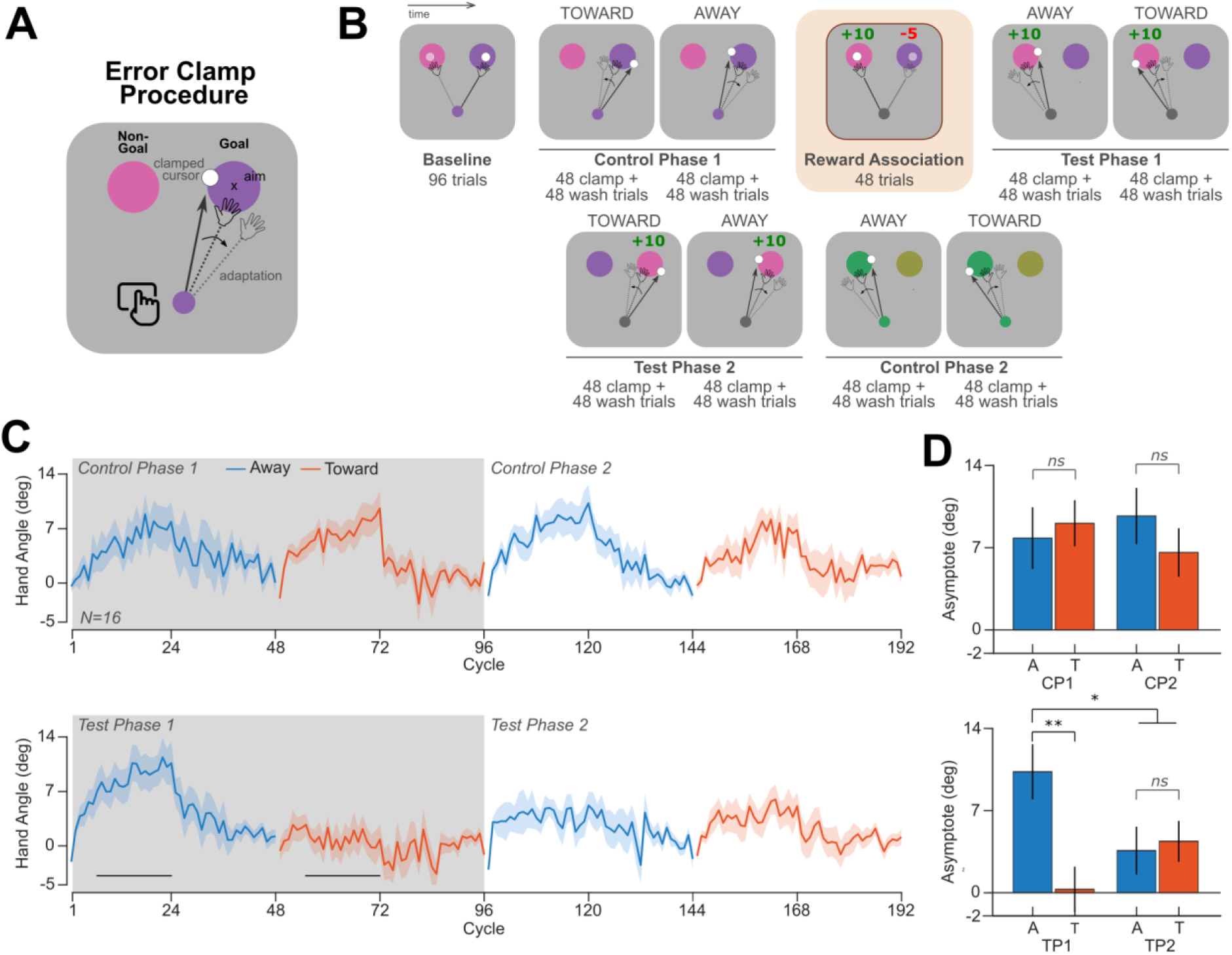
Implicit adaptation is repelled by and suppressed around previously punished movement directions. **A)** General error clamp procedure. Using a trackpad, participants made swiping motions toward one of two visually presented targets, the “goal” and “non-goal” targets. The “x” depicts where participants were instructed to aim, and where they initially move to. They see a fixed, or “clamped”, cursor that creates a sensory-prediction error (a discrepancy between the aiming location and the observed cursor) that drives adaptation in the opposite direction of the observed error. **B)** Task protocol in Experiment 1. After a baseline phase, participants adapted to one of two movement targets in Control Phase 1. The sign of the observed sensorimotor error – given by a fixed (“clamped”) rotational perturbation of visual feedback – alternated across blocks such that the hand either adapted TOWARD or AWAY from the non-goal target. Participants then learned to associate rewards and punishments with each of the two targets. Subsequently, they reached only to the higher-value target, once again with the sign of the error-clamp feedback alternating to produce adaptation TOWARD or AWAY from the low-value target. In Test Phase 2, the target configuration switched; participants still reached to the high-value target but it was now located where the low-value target had been located previously. Finally, target colors changed to neutral colors during Control Phase 2, where participants adapted at the target location opposite the one used in the first Control Phase. **B)** Learning curves (cycle = 2 trials) for Control Phases 1 and 2 (*top*) and Test Phases 1 and 2 (*bottom*). In each learning direction within each phase, the error-clamp was on for 24 cycles and off for the next 24 cycles (washout, see Methods). Shaded regions reflect standard error of the mean. Bars reflect epochs showing significant differences between TOWARD and AWAY learning curves within a phase, as determined by cluster permutation tests (p<0.05). **C)** Adaptation asymptotes for the Control (*top*) and Test Phases (*bottom*). Error bars reflect standard error of the mean. * p<0.05, ** p<0.01.

Overall, participants’ movement data exhibited significant motor learning in all adaptation phases of Experiment 1 (**Fig. 1C**). Adaptation, quantified as the average baseline-corrected hand angle in the last two cycles of the block, averaged 8.4 ± 1.2° and 7.8 ± 1.4° for AWAY and TOWARD blocks, respectively (mean ± SEM) in both Control Phases (**Fig. 1B**). When averaging across both Test Phases 1 and 2, adaptation reached 7.2 ± 1.7° and 2.4 ± 1.3° for AWAY and TOWARD blocks, respectively. Crucially, there was a significant difference between adaptation AWAY and TOWARD when averaging across both Test Phases (4.8°, 95% CI [1.0°, 8.6°], t(15)=2.7, p=0.02, d_z_=0.67), but no such difference between AWAY and TOWARD in the Control Phases (t(15)=0.3, p=0.76). Moreover, the AWAY versus TOWARD differences were significantly larger in the Test Phases than in the Control Phases (t(15)=-2.2, p=0.047, d=0.54). These data illustrate that implicit adaptation appeared to be attenuated when it moved the hand toward a low-value goal, and this effect was not a reflection of baseline motor biases nor due to the mere presence of multiple movement goals in the visual field. Thus, low-value goals appeared to “repel” the effector during adaptation.

After observing that, on average over both Test Phases, adaptation was significantly attenuated when it moved the effector toward the low-value goal, we next asked if this effect was present in both Test Phases individually, or specific to only one Test Phase. If the effect was present in both Test Phases, it would suggest that adaptation is attenuated when it moved the effector toward low-value targets regardless of where they are presented in space (target-identity-based effect). In contrast, if the effect was only present in Test Phase 1, it would suggest that the effect is linked to specific movement directions.

A two-way ANOVA on adaptation over error-clamp direction (TOWARD vs AWAY) and Test Phase (1 vs 2) revealed a significant interaction (F(1,44.4)=8.9, p=0.005, η^2^_partial_ = 0.17), indicating that the effect of the error-clamp direction differed across phases (**Fig. 1D**, bottom). Specifically, adaptation was significantly depressed toward the punished target in Test Phase 1, showing a striking ∼10.1° reduction in asymptotic adaptation in the TOWARD condition versus the AWAY condition (95% CI [3.9, 16.4], t(15)=3.5, p=0.004, d_z_=0.86, **Fig. 1D**); in fact, adaptation toward the punished target in that phase was not significantly different from zero (0.29°, t(15)=0.15, p=0.88). Using cluster permutation tests, we found that the adaptation learning curves in Test Phase 1 significantly diverged as early as cycle 7, and remained significantly different throughout the remainder of the adaptation phase (t_sum_=50.0, p=0.004, **Fig. 1C**). Moreover, 13 out of 16 participants showed reductions in adaptation in the TOWARD versus AWAY block. In contrast, there was no difference in adaptation toward or away from the low-value target in Test Phase 2 (t(14)=-0.16, p=0.87, d_z_=-0.04; *no significant clusters,* **Fig. 1C**). Finally, there was no difference in the effect of TOWARD vs AWAY across the two Control Phases (interaction: F(1,60)=0.9, p=0.3, η^2^_partial_=0.02) and no clusters significantly diverged between TOWARD and AWAY conditions in either Control Phase 1 or 2. These Control Phase results further support the conclusion that our observations in the Test Phases were not driven by baseline motor biases or the visual presence of competing reach goals.

It is important to emphasize that, in Test Phase 2, the rewarded target was now placed where the punished target had been previously located, and thus participants were reaching in a direction that had most recently been punished (**Fig. 1B**). Because participants still only reached to the higher-value target, all movements in Test Phase 2 (in both TOWARD and AWAY blocks) were therefore in the vicinity of a previously punished reach direction. Strikingly, in Test Phase 2 adaptation both TOWARD *and* AWAY from the now-relocated punished target was *suppressed* and not significantly different across the two directions (t(14)=-0.16, p=0.87, d=-0.04, **Fig. 1D**). To quantify this suppression effect, we compared adaptation in Test Phase 2 to the robust adaptation observed in the AWAY block in Test Phase 1 (TOWARD: 6.0° suppression, 95% CI [-0.02, 12.0], t(15)=2.1, p=0.051, d_z_=0.53; AWAY: 7.6° suppression, 95% CI [1.7, 13.5], t(14)=2.8, p=0.02, d_z_=0.71, AVG: 7.4° suppression, 95% CI [2.0, 12.9], t(14)=2.9, p=0.01, d_z_=0.76, **Fig. 1D**). These data suggest that in addition to adaptation being attenuated when it moves the effector toward a low-value reach target, adaptation may also in general be attenuated for reach directions that overlap with previously punished actions.

Further analysis showed that movement durations and reaction times were not significantly different between TOWARD and AWAY blocks in the Control and Test Phases (see Supplemental Table S1). During the Reward Association block, there were no significant differences in movement duration (t(15)=1.1, p=0.29) nor reaction time (t(15)= −1.34, p=0.20) between rewarded and punished trials, nor on trials where participants reached to the rewarded versus punished target (MT: t(15)=-0.69, p=0.50; RT: t(15)=1.42, p=0.18). Finally, in the baseline trials, participants were not significantly biased toward or away from the distractor target when both targets were presented simultaneously (t(15)=0.40, p=0.69, d=0.1); on baseline trials where both targets were presented, reaches deviated from the center of the target by an average of 0.32 ± 0.79° (mean ± SEM), indicating that participants adhered to the instruction to aim for the middle of the cued target.

### Computational modeling of value-modulation of implicit adaptation

The results of Experiment 1 revealed two key patterns in how the reward value of reach goals influenced adaptation. In Test Phase 1, adaptation appeared to be “repelled” away from low-value reach directions, with adaptation away from the low-value target being greater than adaptation toward it. In Test Phase 2, adaptation appeared to be universally “suppressed” around reach directions that had been previously punished.

To explain this pattern of results, we adopted a computational model that was inspired by policy gradient models in reinforcement learning (see Supplemental Note for a theoretical motivation of this model). In our Policy Gradient model (**Fig. 2A**), the agent learns the discrete values of each target and uses those values to inform the weighting of each reach direction. That is, after each trial, the reward feedback obtained on that trial increases (for reward) or decreases (for punishment) the value of that trial’s actual reach direction and also other nearby reach directions, with a Gaussian decay for increasingly dissimilar reach directions. This allows the agent to learn a “value landscape” over all possible reach directions that is high for recently rewarded reaches and low for recently punished reaches. Thus, learning at one location spreads to other similar reach directions. We then assume that this learned value landscape modulates the motor learning rate (of a classic state-space model of implicit motor adaptation^28^. Specifically, the value of the value landscape at a given reach direction determined the learning rate, with low-value reach directions resulting in a diminished learning rate (see Methods). We additionally analyzed models that used this value landscape to modulate the retention parameter or both the learning rate and retention parameters (see Supplemental Note and Fig. S4); the models that only modulated the learning rate best fit the data.

**Figure 2.**
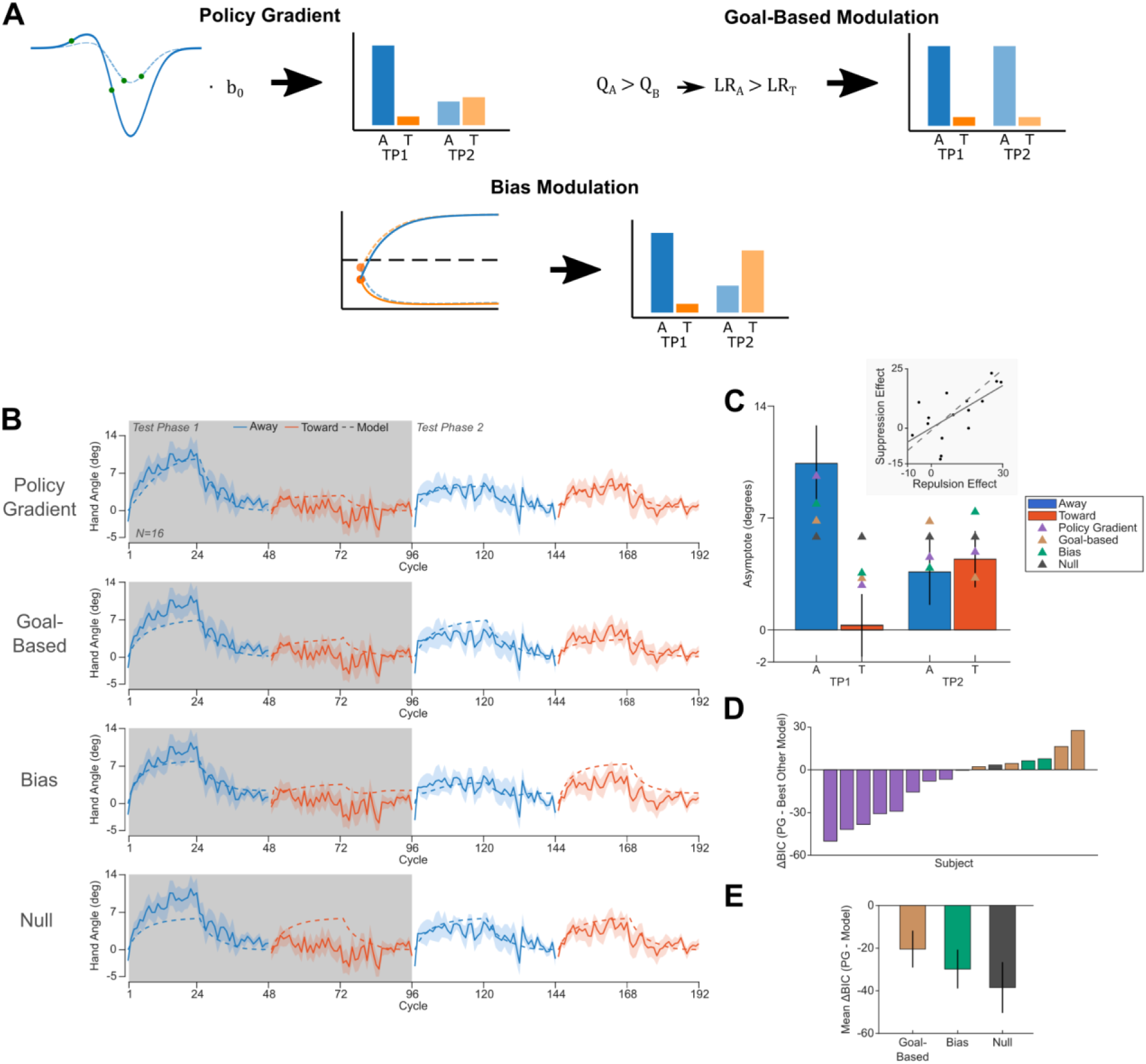
Computational modeling results for Experiment 1. **A)** Schematics of each of the models (not including the Null model). All three models predict repulsion effects (i.e., attenuated adaptation when learning brings the hand towards a punished target) during Test Phase 1, but differ in their predictions for Test Phase 2. Only the Policy Gradient model predicts global suppression effects in Test Phase 2. **B)** Model fits for the Test Phases of Experiment 1 across all four models. Shaded regions reflect standard error of the mean. **C)** *Main*: Model-predicted asymptotes for all four models overlaid over the subject data. Error bars reflect standard error of the mean. *Inset:* Correlation of each subject’s repulsion effect in Test Phase 1 versus their suppression effect in Test Phase 2 (r=0.64, p=0.007). Dashed line reflects the predicted correlation according to the Policy Gradient model. **D)** BIC differences between the Policy Gradient model and the best-performing alternative model for each subject, sorted least to greatest. Lower values reflect better fitting of the Policy Gradient model. Nine of sixteen participants were best fit by the Policy Gradient model, more than double the number seen for the next best (Goal-Based) model. **E)** Mean BIC differences between the Policy Gradient (PG) model and each of the alternative models. Error bars reflect standard error of the mean.

The Policy Gradient model has 5 tunable parameters. The first two, *ϕ_1_* and *ϕ_2_*, correspond to the amplitudes of the Gaussian functions used in the value landscape to represent the values of the high-value and low-value targets respectively. The ratio *λ* = |*ϕ_2_*/*ϕ_1_*| defined the degree of “loss aversion”, indicating how much more aversive the low value target was than the high-value target was appetitive. Together, these two parameters defined the value landscape for all possible reaches; namely, the landscape was the sum of two Gaussians centered at each of the two target locations, with a standard deviation of 20 degrees (matching the target separation; models were fit with the Gaussian widths being a free-parameter but were not justified by BIC, see Supplemental Note and Fig. S5). The next two parameters were *b_0_* and *a* the (baseline) learning rate and retention parameters, respectively, for a state-space model of implicit adaptation (the effective learning rate on each trial was derived from the modulation of the baseline learning rate *b_0_* multiplied by the value of the value landscape at that reach direction). Importantly, the value landscape could never modulate the learning rate to a value greater than 1, the theoretical maximum learning rate. Finally, we introduced a trial-by-trial decay of the value landscape *γ*, which decayed the weights back to 0 over time.

We compared the predictions of our Policy Gradient model to three alternative models. The first alternative model modulated the learning rate of adaptation purely based on the reward value of each of the targets (the “Goal-Based Modulation” model, **Fig. 2A**). This model would predict a lower learning rate for reaches toward a low-valued target (a “repellent” effect) and a higher learning rate for reaches toward a high-value target (an “attraction” effect). This model should not predict significant suppression effects and instead would predict an identical repellent effect to Test Phase 1 in Test Phase 2. This model had 3 parameters: *b_T_* (the learning rate on TOWARD blocks), *b_A_* (the learning rate on AWAY blocks), and *a* (the retention parameter).

Our second alternative model, the “Bias Modulation” model (**Fig. 2A**), assumes that our effect is due to a (hidden) baseline “bias” at the start of each block. In this model, learning proceeds under the usual state-space regime, but instead of hand angles starting at zero (i.e. *x*(0) = 0), there exists an unseen “bias” at the beginning of adaptation (*x*(0) = *β*). In TOWARD blocks, this bias is positive, thus participants begin close to asymptote and have little further to learn. In AWAY blocks, the bias is negative, so participants are farther from asymptote and appear to learn more. This bias is effectively a flat clockwise or counterclockwise shift induced by the Reward Association phase to skew adaptation away from the punished direction. This model has 4 tunable parameters: *a* and *b* for the state space, *β* for the angular bias, and *γ* a trial-by-trial decay of the bias across blocks.

Our third alternative model, the “Null” model, was a benchmark model that assumes that there was no modulation of adaptation by reward history. This model requires only two parameters: *a* and *b*. Adaptation for all blocks was modeled by a single state-space model that used these two parameters. Importantly, aside from the Null model, all models predict the repellent effect observed in Experiment 1 Test Phase 1, but they differ in their predictions for Test Phase 2 (**Fig. 2A**).

### A Policy Gradient model best explains modulation of adaptation by previous reward history in Experiment 1

All four models were fit to each participant’s full adaptation learning curves (including washout) for both the Control and Test Phases. Models were fit with the objective of minimizing the root mean square error (RMSE) between the model-predicted hand angles and subjects’ true hand angles.

Participants’ learning curves were best fit by the Policy Gradient model, with an average RMSE of 6.7° (SD: 1.8°). Out of 16 subjects, 9 were best fit by the Policy Gradient model, 4 by the Goal-Based Modulation model, 2 by the Bias Modulation model, and 1 by the Null model (**Fig. 2D**). The policy gradient model outperformed all three alternative models by BIC (mean ΔBIC relative to the Policy Gradient model; Goal-based: 19.8; Bias: 29.4; Null: 37.4, **Fig. 2E**). Figure 2B shows the model predictions for Test Phases 1 and 2 for each of the models. The Policy Gradient model most closely predicted the asymptotes for the both TOWARD and AWAY blocks of Test Phases 1 and 2, with an average error of 0.7° (SD: 5.5°) (TP1 A: −0.8 ± 4.0°; TP1 T: 2.5 ± 6.6°; TP2 A: 0.6 ± 5.7°; TP2 T: 0.5 ± 5.3°) (**Fig. 2C**).

While all alternative models predicted the repellent effect observed in Test Phase 1, the Policy Gradient model best fit the suppression effect observed in Test Phase 2, and also better explained variance in the learning curves across both Control and Test Phases (Control Phase fits are shown in Fig. S2). Further, as predicted by the Policy Gradient model, each subject’s degree of repulsion in Test Phase 1 correlated strongly with their degree of suppression in Test Phase 2 (Pearson correlation: r=0.64, t(14)=3.17, p=0.007, **Fig. 2C** *inset*). This provides robust evidence for the validity of the Policy Gradient model in explaining the pattern of results observed in Experiment 1. We next performed additional experiments to further test how the learned value of particular reach directions may influence implicit adaptation in accordance with a Policy Gradient model.

### Suppression of adaptation toward low-value goals is tied to reach direction and recent reinforcement history

The results of Experiment 1 demonstrated that implicit adaptation is attenuated when adaptation brings the hand toward a low-valued (negatively-valued) goal, reflecting a kind of repelling effect. This effect went away when the target locations were swapped: In Test Phase 2 (where the rewarded target was placed in the location where participants were previously punished), adaptation in both directions was globally suppressed. This finding suggests that adaptation is suppressed in the vicinity of previously punished reach directions, whether or not those directions reflect the current movement goal or are simply near the current goal. In other words, a history of previously punished actions locally suppresses adaptation for nearby movements. Our Policy Gradient model accounts for this by separately modeling the discrete value of the targets from the continuous value of all possible reach directions (akin to an “actor-critic” reinforcement learning agent, see Supplemental Note). In the Policy Gradient model, the value landscape is learned directly, on a trial-by-trial basis, from the recent reinforcement history for each reach direction, with learning at one location spreading to other nearby reach directions.

To further test the model, in Experiment 2 we added a second Reward Association “top-up” block following Test Phase 1, with the target configurations switched such that the Reward Association top-up block ended with the punished and rewarded targets in the same configuration as they would be presented in Test Phase 2 (**Fig. 3A**). That is, this manipulation ensured that in Test Phase 2 participants had most recently experienced punishment at the *new* location of the punished target, and had most recently experienced reward at the new location of the rewarded target. Thus, by undoing the learned association of punishment with the new location of the rewarded target, and by establishing an aversive association with the new location of the punished target, we expected to “rescue” the directionally-specific repelling effect (TOWARD vs AWAY difference) that we observed in Test Phase 1 of Experiment 1. We thus expected to now see repelling effects in both Test Phases.

**Figure 3.**
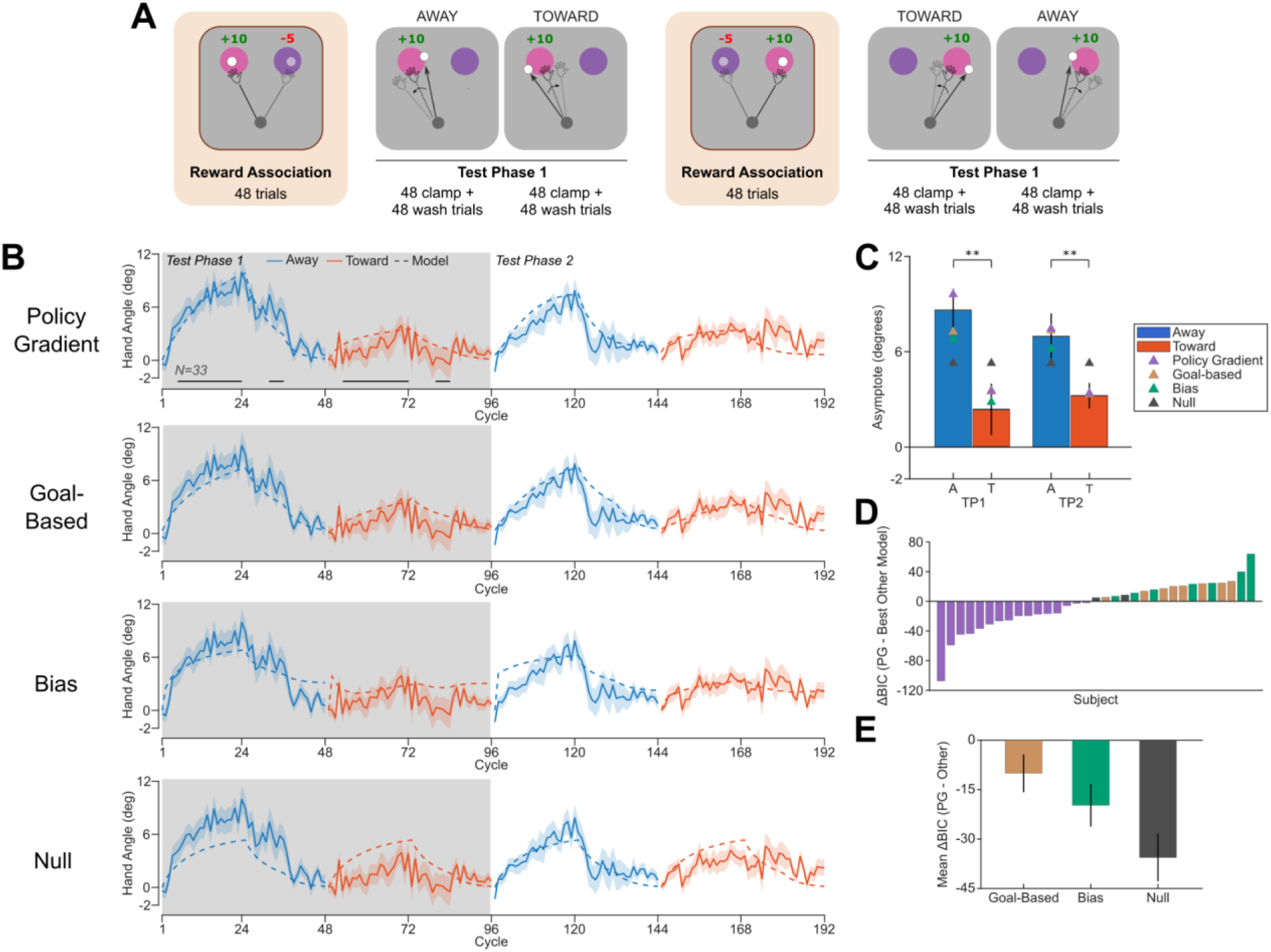
Experiment 2 behavior and modeling results. **A)** Experiment 2 Task protocol. Test Phases were identical to Experiment 1, except that the Reward Association block was repeated between Test Phase 1 and 2, this time with the target configurations swapped (*see Methods*). **B)** Model fits for both Test Phases. Black bars in the top row reflect epochs with significant differences between TOWARD and AWAY adaptation directions, via cluster permutation tests (p<0.05). Shaded regions reflect standard error of the mean. **C)** Asymptotes for both Test Phases, with model-predicted asymptotes overlaid. Error bars reflect standard error of the mean. **D)** BIC differences between the Policy Gradient model and the best alternative model. Sixteen of thirty-three participants were best fit by the Policy Gradient model, twice that seen for the next best model. **E)** Mean BIC differences between the Policy Gradient (PG) model and each of the alternative models. Error bars reflect standard error of the mean. * p<0.05 ** p<0.01.

We replicated the repelling effects in Test Phase 1: Adaptation was significantly suppressed in the TOWARD versus AWAY conditions, by an average of 7.1° (95% CI [2.5, 11.8], t(31)=3.1, p=0.004, d_z_=0.55, **Fig. 3B, C**). Cluster permutation tests showed that adaptation learning curves significantly diverged as early as cycle 5 and remained significantly different throughout adaptation (largest cluster: t_sum_=59.5, p=0.0003, **Fig. 3B**). As in Experiment 1, adaptation TOWARD in Test Phase 1 was not significantly different from zero (mean: 2.3°, 95% CI [-1.1, 5.7], t(31)=1.4, p=0.18, **Fig. 3C**).

The key question in Experiment 2 was if adaptation was significantly different across directions (TOWARD vs AWAY) in Test Phase 2, which followed the new Reward Association top-up block. Indeed, adaptation did significantly differ across learning directions (TOWARD>AWAY, 3.9°, 95% CI [1.2, 6.7], t(31)=2.9, p=0.007, d_z_=0.51, **Fig. 3C**), though we note that the more conservative exploratory cluster permutation tests across individual trial cycles failed to identify significant epochs after correcting for multiple comparisons (**Fig. 3B**). These findings show that, as hypothesized, recent reinforcement history drives the observed repulsion effect of punished movements.

Moreover, the subtly weaker dissociation between the TOWARD and AWAY curves we observed in Test Phase 2 versus Test Phase 1 (**Fig. 3C**) was also predicted by the Policy Gradient model. This is because the new value landscape learned in the top-up reward association block must be learned on top of the original value landscape, resulting in interference between the two value gradients as previously punished reach directions are now rewarded. The Goal-Based Modulation model does not account for this interference, as learning rates are solely modulated by the value of the targets themselves.

As in Experiment 1, the Policy Gradient model best fit the data in Experiment 2, achieving an average RMSE of 5.5° (SD: 2.3°). This is despite the fact that the suppression effects in Experiment 1 – which provide the strongest modeling distinctions – were abolished in Experiment 2. Out of 33 participants, 16 were best fit by the Policy Gradient model, 8 by the Goal-Based Modulation model, 7 by the Bias Modulation model, and 2 by the Null model (**Fig. 3D)**. BIC comparison provided robust evidence that the Policy Gradient model best fit the subject data on average (mean ΔBIC relative to the Policy Gradient model; Goal-Based: 10.0; Bias: 19.8; Null: 35.6; **Fig. 3E**). The Policy Gradient model most closely predicted the asymptotes for both TOWARD and AWAY blocks of Test Phases 1 and 2, with an average error of 0.7° (SD: 5.6°) (mean ± SD, TP1 A: 1.0 ± 4.6°; TP1 T: 1.1 ± 8.5°; TP2 A: 0.1 ± 4.8°; TP2 T: 0.1 ± 3.8°, **Fig. 3C**).

### Adaptation is suppressed toward relatively low-value goals but not enhanced toward high-value goals

Our results thus far demonstrate that reinforcement history can attenuate adaptation in the vicinity of previously punished reach directions, both when adaptation moves the hand toward those targets even when they are not the current goal (repelling effects), and when the punished movement direction is itself the current goal (suppression effects). Mirroring these effects, we reasoned that *rewarding* particular reach directions could locally enhance adaptation as well. In Experiment 3, we made the lower-valued target neutral (+0) so that no actions were punished throughout the experiment. Participants learned a modified reward association such that the rewarded target always provided +10 points and the neutral target always provided +0 points (**Fig. 4A**). After learning the association, participants were instructed to always reach to the neutral target and adaptation was induced using error-clamped perturbation feedback, with an alternating sign such that adaptation would bring participants’ reaches either TOWARD or AWAY from the *rewarded* target. Mirroring Experiment 1, in Test Phase 1 of Experiment 3 the target configuration matched the final configuration of the Reward Association block; in Test Phase 2 the configuration swapped such that the neutral target was placed where the rewarded target was located previously (**Fig. 4A**).

**Figure 4.**
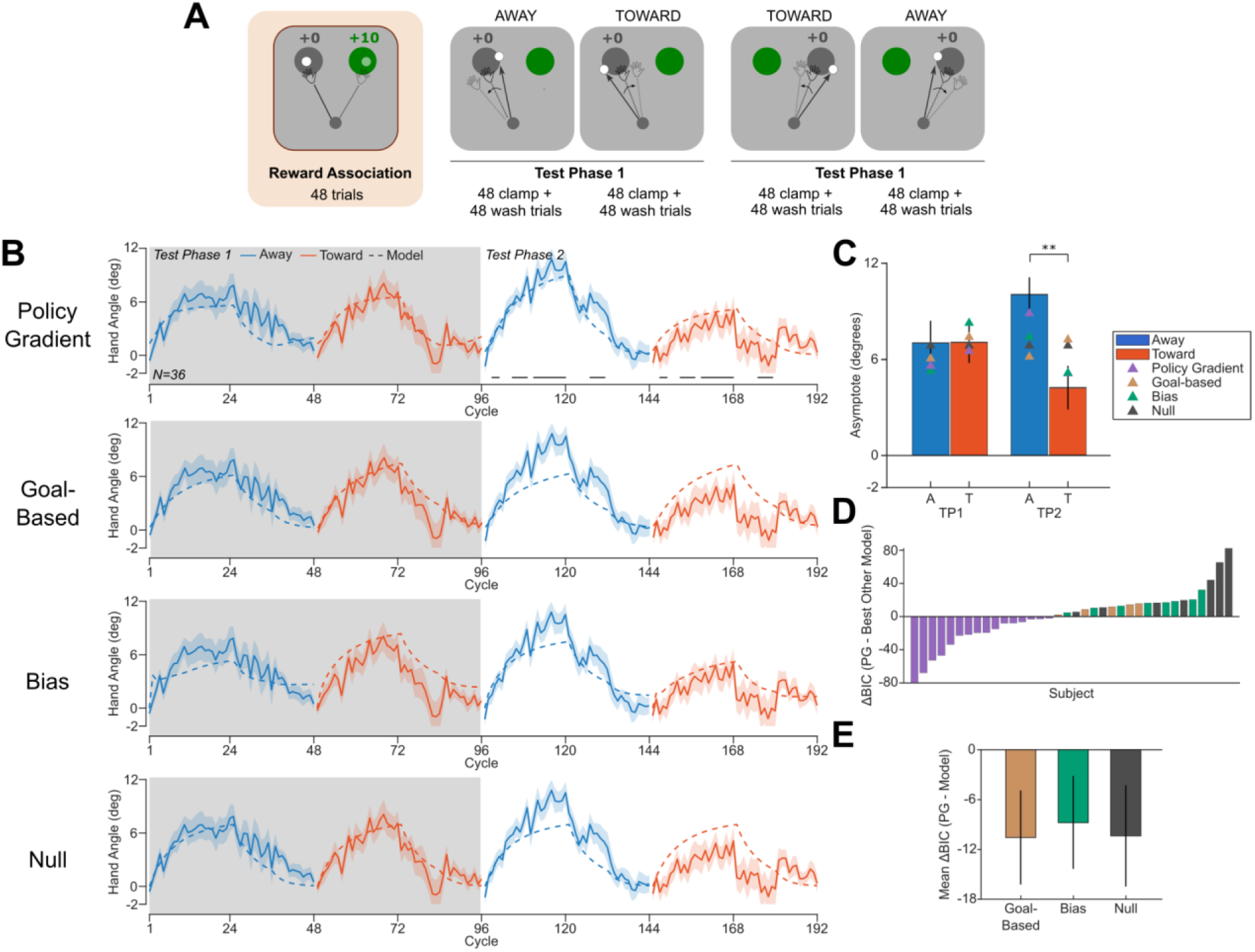
Experiment 3 behavior and modeling results. **A)** Experiment 3 Task protocol. The lower-value target was now neutral (+0 reward) and participants adapted around the neutral target in both Test Phases (*see Methods*). **B** Model fits for both Test Phases. Black bars in the top row reflect epochs with significant differences between TOWARD and AWAY adaptation directions, via cluster permutation tests (p<0.05). Shaded regions reflect standard error of the mean. **C)** Asymptotes for both Test Phases, with model-predicted asymptotes overlaid. Error bars reflect standard error of the mean. **D)** BIC differences between the Policy Gradient model and the best alternative model. Sixteen of thirty-six participants were best fit by the Policy Gradient model, twice that seen for the next best model. **E)** Mean BIC differences between the Policy Gradient (PG) model and each of the alternative models. Error bars reflect standard error of the mean. * p<0.05 ** p<0.01.

In Test Phase 1 there was no significant difference between adaptation TOWARD versus AWAY from the rewarded target (t(33)=-0.17, p=0.87, **Fig. 4C**), nor were any significant divergences in the learning curve identified via cluster permutation tests (**Fig. 4B**). This suggests that, at least with the values we used, there is not a robust “attraction” effect when adaptation brings the hand toward high-value targets. Interestingly, in Test Phase 1 both adaptation directions exhibited suppressed adaptation relative to the AWAY block in Test Phase 2 (TOWARD: 2.9° suppression, 95% CI [0.4, 5.4], t(33)=2.4, p=0.02, d_z_=0.40; AWAY: 3.2° suppression, 95% CI [0.3, 6.1], t(32)=2.2, p=0.03, d_z_=0.38). These results suggest that despite the neutral value of the goal target in the Test Phases, its *relative* devaluation compared to the rewarded target appeared to suppress adaptation in both directions, mirroring the Test Phase 2 results of Experiment 1.

Moreover, adaptation was significantly different between error-clamp directions in Test Phase 2, when the neutral target (the goal target) was now placed where the rewarded target had been located previously. That is, adaptation was now significantly attenuated when it moved hand toward the rewarded target identity/color, which was located where the lower-valued neutral target had been previously) (6.1° reduction, 95% CI [2.5, 9.7], t(34)=3.5, p=0.002, d_z_=0.58, **Fig. 3D**), with multiple epochs showing significant divergences in the learning curve (largest cluster: t_sum_=31.7, p=0.002, **Fig. 3C** *right*). A two-way ANOVA on adaptation over clamp direction (TOWARD vs AWAY) and phase (1 vs 2) produced a significant interaction term (F(1, 102.8)=6.8, p=0.01), indicating that the effect of clamp direction differed across Test Phases. In other words, Experiment 3 Test Phase 2 recapitulated the repelling effects of Experiment 1 Test Phase 1, but now adaptation was repelled away from a relatively devalued *neutral* reach direction.

These results support a surprising and informative extension of Experiments 1 and 2: Motor adaptation is suppressed when it brings the effector toward *relatively* low-valued reach directions, not merely *punished* (negatively-valued) reach directions. In other words, modulation of adaptation by reinforcement history appears to be sensitive to the learned relative values rather than the absolute values of movements. This pattern of results was successfully predicted by the Policy Gradient model, but not by any other model. Specifically, the Goal-Based Modulation and Bias Modulation models predicted a strong “attraction” effect in Experiment 3 Test Phase 1 (i.e., that learning would be greater toward the high-value target than away from it), which was not present. The Goal-Based Modulation model predicts the same “attraction” effect in Test Phase 2, which is the opposite of the observed repelling effect. While the Policy Gradient model does numerically predict a small attraction effect, the effect is quite weak, because reward modulation cannot make the effective learning rate greater than 1 and the high degree of loss aversion drives the effect of punishment to dominate over the effect of reward.

The Policy Gradient once again best fit the data in Experiment 3, with an average RMSE of 6.5° (SD: 2.0°). Out of 36 participants, 16 were best fit by the Policy Gradient model, 5 by the Goal-Based model, 8 by the Bias model, and 7 by the Null model (**Fig. 4D**). BIC comparison provided further evidence that the Policy Gradient model best fit the subject data on average (mean ΔBIC relative to the Policy Gradient model; Goal-Based: 10.6; Bias: 8.7; Null: 10.4; **Fig. 4E**). The Policy Gradient model most closely predicted the asymptotes for both TOWARD and AWAY blocks of Test Phases 1 and 2, with an average error of −0.5° (SD: 5.5°) (mean ± SD, TP1 A: −1.4 ± 5.6°; TP1 T: −0.6 ± 4.8°; TP2 A: −1.1 ± 4.7°; TP2 T: 0.9 ± 6.6°, **Fig. 4C**).

### Reward-learning and adaptation-related parameters were preserved across experiments

Across our three experiments, the Policy Gradient model was the best fitting model, despite all three experiments having disparate reward contexts and block designs. Moreover, Experiments 2 and 3 were both conducted online using crowd-sourced participants using their own devices, while Experiment 1 was conducted in-lab using a custom trackpad. Experiments 1 and 2 had identical Reward Association blocks which differed greatly from the Reward Association block in Experiment 3, where outcomes were deterministic (not probabilistic, see *Methods*) and the lower-value target was neutral, instead of punished. Despite these significant methodological differences across experiments, the parameters of the Policy Gradient model were remarkably conserved (**Fig. 5**).

**Figure 5.**
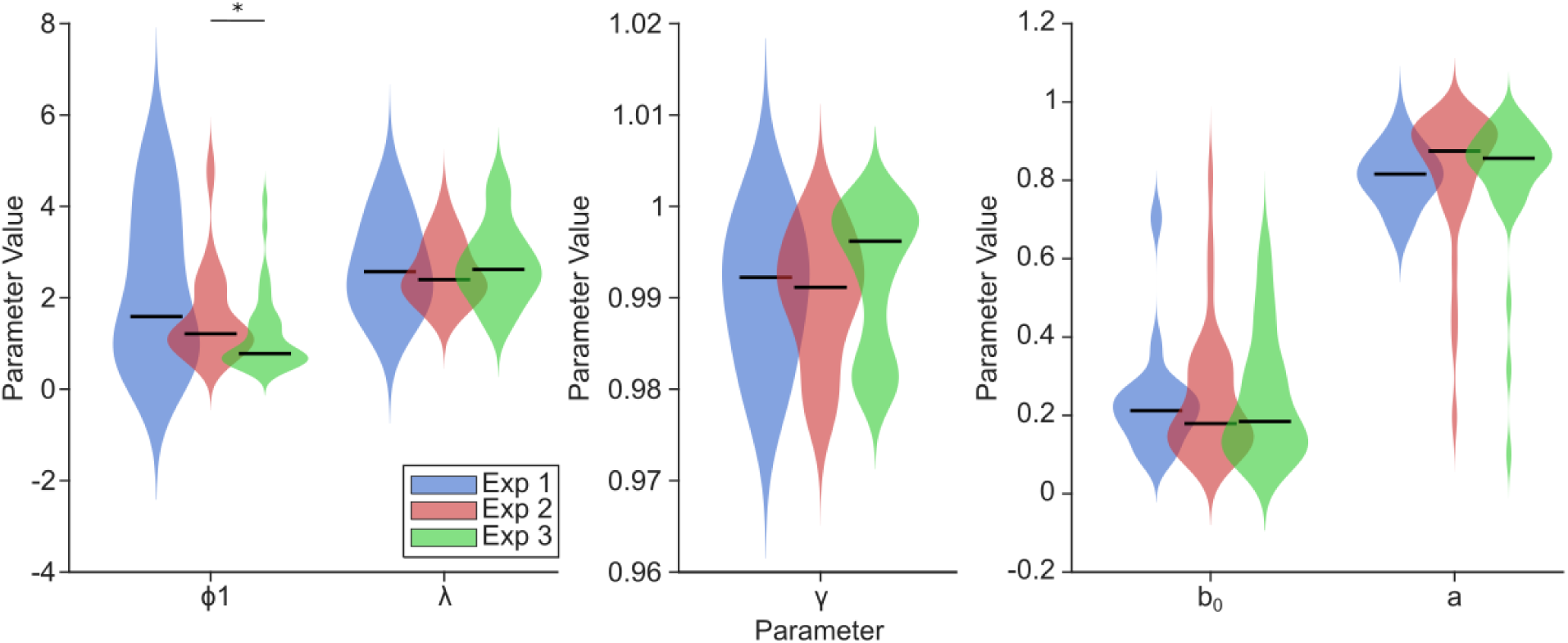
Policy Gradient model parameter comparison across Exps. 1-3. *Left:* Reward-related parameters, reward amplitude Φ_1_ and loss aversion λ. *Middle:* landscape forgetting parameter γ. *Right:* Adaptation-related parameters, baseline learning rate b_0_ and retention a. Parameters showed preservation across all three experiments, acting as a strong validity check. Capacity for adaptation was comparable across all three experiments. Reward amplitudes were significantly different between Exps 2 and 3, likely due to differences in the Reward Association learned in each experiment. * p<0.05.

In Experiments 1 and 2, where the Reward Association blocks were identical, the reward amplitude (U=425, p=0.60, |r|=0.07, BF_01_=1.3) and loss aversion (U=413, p=0.79, |r|=0.04, BF_01_= 2.9) parameters were not significantly different. This suggests that, across the population, reward learning was similar for both groups of participants despite one experiment being conducted online and the other in-person. In fact, the loss aversion parameters were strikingly consistent with values obtained in nonmotor economic decision-making tasks, averaging ∼2.7^29^. However, the reward amplitudes were subtly but significantly different between Experiment 2 and Experiment 3 (U=1348, p=0.02, |r|=0.28) and marginally significant between Experiments 1 and 3 (U=515, p=0.07, |r|=0.25, BF_10_=7.8). These results can likely be explained by the fact that Experiment 3 had a distinct Reward Association block, so participants learned a different value landscape in this experiment compared to the other two.

The degree to which the learned value gradient passively decayed (back to being flat) over time was also preserved across all three experiments (Exp. 1 vs Exp. 2: U=423, p=0.63, |r|=0.07, BF_01_=3.3; Exp. 1 vs Exp. 3: U=368, p=0.27, |r|=0.15, BF_01_=2.8; Exp. 2 vs Exp. 3: U=1017, p=0.10, |r|=0.20, BF_01_=2.3). Similarly, the adaptation-related parameters *b_0_* and *a* were not significantly different across the three experiments (*b_0_*, Exp. 1 vs Exp. 2: U=408, p=0.87, |r|=0.02, BF_01_=3.3; Exp. 1 vs Exp. 3: U=429, p=0.93, |r|=0.01, BF_01_=3.3; Exp. 2 vs Exp. 3: U=1191, p=0.67, |r|=0.05, BF_01_=4.0; *a*, Exp. 1 vs Exp. 2: U=323, p=0.10, |r|=0.23, BF_01_=3.3, Exp. 1 vs Exp. 3: U=358, p=0.19, |r|=0.18, BF_01_=3.4, Exp. 2 vs Exp. 3: U=1234, p=0.35, |r|=0.11, BF_01_=3.9). This suggests that the three groups of participants largely demonstrated the same intrinsic capacity for implicit adaptation, further validating our modeling approach.

## Discussion

How do the values of actions inform how we unconsciously modify them? Recent studies of human motor learning point to an interaction between value-based learning and error-based implicit motor adaptation^6,8,9^. However, the interpretation of these results has been complicated by various alternative explanations, including the use of explicit strategies and task error signals associated with non-adaptation-based forms of motor learning^10,11,18^. Here, we took a novel approach to resolving this controversy by first training participants on the relative values of two competing action goals in the environment, and then asking if this learned association shapes implicit adaptation in the vicinity of these goals. Across three experiments, we demonstrate that implicit adaptation is attenuated around low-value reach directions (**Figs. 2-4**). These findings point to a close interaction between value-based learning and implicit motor adaptation.

We suggest that reinforcement learning influences implicit adaptation by influencing the value of particular reach directions, as might be predicted by policy-gradient RL models^30,31^. In our proposal, when a reach direction is punished, the error sensitivity for reaches in that direction, and other similar directions, is decreased. In other words, adaptation is suppressed for reaches that are kinematically similar to that punished direction, effectively diminishing updates to the agent’s action policy that might lead to movements similar to ones that were recently punished. This interaction would serve a clear adaptive purpose: by preventing large updates in the direction of a previously punished action, the agent may avoid reproducing a potentially maladaptive action when trying to account for motor errors.

We highlight that only our Policy Gradient model could explain both the repelling effects observed across experiments and the robust suppression effects observed in Test Phase 2 of Experiment 1. These results can be simply summarized by stating that adaptation is suppressed around reach directions that have been previously punished. However, it is important to note that the high-value target in Test Phase 1 was located 20 degrees away from the low-valued target. Thus, the suppression of adaptation in the TOWARD blocks occurs when the hand is a full 20 degrees *away* from the low-valued target. How is the low-valued target able to exert such a strong influence at such a distance? And how is the suppression in Test Phase 1 stronger at 20 degrees away than it is when the effector is reaching directly toward those low-valued directions in Test Phase 2? Only a model that explicitly accounts for the reward value of all possible reach directions, including reach directions that the participant never reached to, can account for these results, as our Policy Gradient model does (**Fig. 2A**).

In Experiment 2, we extended our findings from Experiment 1 by showing that we could reverse the learned movement-reward association and produce globally suppressed adaptation when errors are experienced when moving toward the most-recently punished reach directions. Importantly, while the Goal-Based Modulation model could predict repulsion away from the low-value target for both Test Phases in Experiment 2 (and with fewer parameters), our Policy Gradient model still best fit the data. This is due to its prediction of interference between the initially learned value landscape and the reversed landscape learned in the second reward association phase.

In addition to adaptation being suppressed in the vicinity of punished reaches, adaptation could in theory be enhanced toward rewarded reaches. However, our data did not support this hypothesis. Not only was adaptation not significantly different between TOWARD and AWAY blocks in Test Phase 2 of Experiment 1 (where an attraction effect could have been present), but we also failed to find a “reward-attraction” effect in Test Phase 1 of Experiment 3. This valence asymmetry could be due to numerous factors. First, there may be an upper limit on the implicit adaptation system’s sensitivity to errors (as implemented in our Policy Gradient model), such that rewards cannot enhance the sensitivity to error beyond this maximum, while punishments can strongly depress error sensitivity and attenuate learning. Second, as captured in our loss-aversion parameter, it may be that the rewards used in our task were simply not as salient as the punishments. Future research using a similar paradigm could test a broader range of reward feedback values to better explain the asymmetry we observed here.

At the neural level, the influence of value-based learning on implicit adaptation could be related to communication between the basal ganglia and cerebellum^32^, a circuit where policy-gradient mechanisms have already been proposed to explain motor behavior^30^. For example, recent neuroimaging work in humans has suggested that reinforcement learning circuits in the basal ganglia and error-based learning circuits of the cerebellum reciprocally inhibit each other, such that reinforcement feedback can inhibit learning from errors^33^. And prior work has highlighted the crucial role of the striatum as the interface between reward and motor performance^34^. Here, we highlight that punishment may similarly locally suppress adaptation to motor errors. This proposed cross-talk between the basal ganglia and cerebellum may serve to modulate motor learning in a manner that ultimately avoids potentially maladaptive or harmful actions. Moreover, the cerebellum itself has been shown to encode reward, punishment, and other reinforcement learning signals like reward prediction errors^35–39^. Thus, intra-cerebellar processes themselves may also underlie interactions between the value domain and motor learning. Further investigations, perhaps using functional neuroimaging, could offer clues about the underlying neural mechanisms of the suppression of adaptation we observed here.

In conclusion, our study provides novel behavioral and computational evidence that implicit motor adaptation is sensitive to value-based information about actions. These findings point to an “Instrumental-Motor Transfer” effect between reinforcement learning computations and motor memories, suggesting a close link between action selection and action execution processes.

## Materials and Methods

### Subjects

Experiment 1: 16 right-handed subjects (6M, 10F, aged 20-36, mean age: 26.8 ± 4.5) were recruited from the Johns Hopkins community to participate in Experiment 1. Written consent was obtained from all participants in accordance with procedures approved by the Johns Hopkins School of Medicine Institutional Review Board (IRB00283000) and participants were compensated with a show-up fee of $25 for the 1.5 hour study, with an additional $10 bonus or $5 penalty for a randomly selected trial during the experiment to motivate them to attend closely to the task (*see Experiment 1 Protocol*). 15 out of 16 subjects received the bonus.

Experiments 2 and 3 were crowd-sourced using the online platform Prolific. A total of 91 participants completed the experiments online (Exp. 2: 47, Exp. 1: 44). Consent was obtained from all participants in accordance with the Yale University Institutional Review Board (Protocol 2000027351). Recruitment was restricted to right-handed or ambidextrous individuals in the United States between the ages of 18 and 35, who had at least 40 prior Prolific submissions. However, 22 participants were excluded due to failure to comply with task instructions (*see Data Analysis*; Exp. 2: 14; Exp. 3: 8). The final sample included 33 participants in Experiment 2 (14M, 19F, aged 22-35, mean age: 29.9 ± 3.6) and 36 participants in Experiment 3 (10M, 26F, aged: 19-35, mean age: 30.0 ± 4.5). Participants were compensated at a rate of $10/hr for the experiments.

### Apparatus

Participants recruited for Experiment 1 completed the task in a mock fMRI scanner (participants were run in the mock scanner in preparation for future study follow-ups using functional neuroimaging). Participants lay flat on a moveable platform. Their responses were recorded on a wired USB computer trackpad (Perixx PERIPAD-506, active area: 5”x4”) connected to the task computer. Task stimuli were displayed on a monitor in the mock scanner and reflected onto a mirror positioned 18 inches away and directly over the participant’s head, angled so that the stimuli on the monitor would be visible to the participant while laying down. Task stimuli and movement recording were handled by custom written MATLAB scripts (MATLAB R2025a, Psychtoolbox 3.0.16) and trackpad movements were recorded at a frequency of 120Hz.

### General Task Procedure

Participants were instructed to make fast but accurate slicing motions using a trackpad (or mouse, if online) toward one of two targets presented on the screen. Two targets (16mm or 60px diameter; Exps. 1-2: pink and purple; Exp. 3: green and gray) were presented 80mm (in-lab) or 300px (online) from the central starting position (6mm or 24px diameter), centered in the middle of the screen. Initially, the goal target was indicated by the color of the central starting location, presented on the screen for 500ms prior to the appearance of the two targets. After learning the reward association to each of the targets (see below), participants were instructed to always go to either the more rewarded (Exps. 1-2) or less rewarded (Exp. 3) target; on these trials, the central starting location would be colored a neutral light blue color so as to not indicate any specific target.

Trials could provide one of three types of feedback. On veridical trials, participants controlled a light blue cursor (4mm or 16px diameter) and veridical feedback was provided throughout the entire reach and at endpoint. The cursor would freeze when the reach amplitude exceeded the target radius (80mm or 300px). On “clamp” trials, the cursor was also displayed throughout the entire reach and terminated at the target radius. However, the trajectory was constrained along a fixed trajectory, 5.5° offset from the center of the goal target, such that the cursor would straddle the edge of the target on every trial^9^ (**Fig. 1A**). Finally, on no-feedback trials, no feedback was provided during movement and the trial ended after the participant’s reach amplitude exceeded the target’s radial distance.

If a trial was rewarded (see below), numerical points were provided only if the displayed cursor (veridical or clamp) landed within the target (i.e., points were provided on all clamp trials). Point feedback was provided at the same angular position as the cursor feedback, at a radial amplitude that exceeded the target distance by 20% so that the text would be visible beyond the target. If the cursor missed the target or was not displayed (as in no-feedback trials), no points were displayed.

Trials would begin with the presentation of the central starting location in the middle of the screen for 500ms. Initially, the color of the starting location indicated which target was the “goal” target. When the starting cue disappeared, it was replaced by the cursor, and the two targets appeared. The pair of targets was presented either at [125° & 145°] or [305° & 325°], separated by 20°. Target locations alternated pseudorandomly such that targets could not appear in the same quadrant for more than 4 trials in a row. Data from both quadrants was rotated to a common reference frame and averaged for all analyses.

Upon the appearance of the targets, participants were instructed to make a quick but accurate swiping motion through the middle of the goal target. If the displayed cursor landed on the target and the trial was rewarded, point feedback was provided along with endpoint feedback. Otherwise, only endpoint feedback was provided. If the participant’s reach was greater than 30° away from the goal target, the reach duration exceeded 300 ms, the reaction time (time from target appearance to reach initiation) exceeded 1000ms, or if the reach trajectory was not straight (median absolute angular deviation greater than 10°), endpoint feedback was replaced (Exps. 2 & 3) or followed by (Exp. 1) a warning screen for 1000ms telling participants what their specific error was (e.g. “Please start the reach sooner” if reaction time exceeded 1000ms). A short reach duration was emphasized to ensure straight ballistic reaches toward the target without feedback corrections.

### Experiment 1 Procedure

All trials in Exp. 1 were constrained to last 3.6 seconds. The central starting location was displayed on the screen for 500ms, and a fixed 800 ms intertrial interval was enforced. The remainder of the 3.6 seconds was divided between reaction time, reach duration, and endpoint feedback. For instance, if a participant had a 500ms reaction time, 100ms reach duration, endpoint feedback was provided for 1700 ms and then the screen would go blank except for a central fixation cross for a 800 ms intertrial interval.

The experiment began with a 96-trial baseline block where the goal target color and location randomly varied. This phase was designed to help participants gain familiarity with the center-out reaching task, and with moving to the target whose color matched the central starting cue. Randomly interleaved throughout the block were 48 trials where both targets were presented and 48 trials where only one target was presented (12 repetitions at each of the 4 possible target locations, 6 repetitions for each color). This was done to measure and control for any baseline biases in reach direction caused by the presence of an alternative target^40^. Target configurations and quadrants alternated pseudorandomly such that no quadrant or configuration would appear more than 4 trials in a row. In total, participants reached equally to each color and target location, reaching to the target whose color matched the color of the central starting location presented at the start of the trial. Participants were instructed to focus on reaching quickly but accurately through the middle of the goal target.

After the baseline phase, participants were informed that the cursor would now be “clamped” to a single direction: it would follow a fixed trajectory terminating slightly off-center on the edge of the target, no matter where they actually moved their hand. Participants were instructed to ignore the cursor and focus on bringing their actual reach to the center of the target, even though they would not receive veridical feedback. This fixed error-clamp task has been shown to reliably induce implicit motor adaptation without inducing explicit re-aiming strategies^27,41,42^. We reasoned that by having the cursor still visibly contact the target, participants would not explicitly interpret the movement as a “miss”, though we still expected to see robust adaptation to the imposed subtle perturbation, consistent with previous work^9^,^18^.

After baseline, participants then began Control Phase 1 (**Fig. 1B**). This consisted of two 48-trial blocks of clamped feedback at one of the two targets, with each clamp block followed by a 48-trial washout block that was designed to de-adapt movements back to baseline. During washout blocks, the first 24 trials were no-feedback, followed by 24 trials of veridical feedback to fully wash out implicit adaptation^41^. Full analysis of the washout blocks is included in the Supplemental Materials (Fig. S1). The two clamp blocks alternated the sign (+/-) of the clamped cursor rotation such that adaptation would proceed either toward the alternative target (TOWARD) or away from it (AWAY). Whether participants started with a TOWARD or AWAY block was counterbalanced. Since Control Phase 1 always went to one target, we counterbalanced the color and location of that target. We note that all adaptation in the Control Phase clamp blocks occurs without any learning about the “values” of the targets or particular reach directions, allowing us to compare later adaptation behavior to the Control Phase data.

After completing the four blocks of Control Phase 1, participants next completed the 48-trial Reward Association Phase. Here the configuration of the targets swapped, such that if the red target was placed clockwise relative to the green target in Control Phase 1, it would now be counterclockwise (and vice versa). The Reward Association Phase used a forced-choice design, where subjects were instructed to reach to a specific target on each trial. When the veridical cursor feedback landed in the instructed target, participants would receive points. One of the two targets was designated the “rewarded target”, which provided rewards (+10 points) on 80% of trials and took away points (−5) on 20% of trials. The rewarded target was always the opposite color as the target participants reached to in Control Phase 1, to ensure that participants were paying attention to the task contingencies rather than adopting a simple strategy of recalling the color of the target they most frequently reached to. The other target, the “punished target”, subtracted points (−5) on 80% of trials and provided points (+10) on 20% of trials. We emphasize that participants were instructed to always reach to the target that matched the color of the central starting location, even if this involved going to the punished target and likely losing points. They were further instructed to track which target provided more points than the other, and were informed that they would be later asked to identify the more favorable target at the end of the block. For the first 12 trials, participants went to one color target. They then went to the other target for the next 12 trials. After these first 24 trials, the target configuration switched again. Participants continued reaching to the same color target, now in a different location, for 12 trials. Finally, for the last 12 trials, participants went to the other color target. This ensured that participants received equal amounts of reward and punishment history at each reach location and that the association was only to the color of the target, and we counterbalanced whether participants started at the rewarded or punished target. To ensure that at least 80% of trials were rewarded at the rewarded target and punished at the punished target, we kept track of the percentage of reversed outcomes (e.g. punishment at the rewarded target) and, if this exceeded 20% within the block, the random outcome was overwritten. At the end of the Reward Association block, participants were asked to explicitly identify the rewarded target. All participants correctly identified the rewarded target.

The final configuration of the targets in the Reward Association Phase was maintained for Test Phase 1 (i.e. if the pink target was clockwise relative to the purple target at the end of the Reward Association block, it would remain that way for Test Phase 1). Just like Control Phase 1, this block consisted of two 48-trial error-clamp blocks, each followed by a 48-trial washout block. Participants were now instructed to always go to the rewarded target, and this instruction was persistently displayed in the upper right corner of the screen on every trial. The central starting location was now colored a neutral light blue color that matched the cursor. The clamped cursor error thus always straddled the edge of the rewarded target. Here again, due to the alternating sign of the clamped cursor, one of the two blocks would adapt movements toward the punished target (TOWARD) or away from it (AWAY). Next followed Test Phase 2, which was identical to Test Phase 1 except that the target configurations switched again, such that the rewarded target was now in the location where the punished target was located previously.

After Test Phase 2, participants completed Control Phase 2. This phase was identical to Control Phase 1 except for two key differences: First, the reach location that participants went to was the opposite of where they went in Control Phase 1 (i.e. if in Control Phase 1, participants reached to the more clockwise target, they now reached to the more counterclockwise target). Additionally, since the original colors now had rewards associated with them, we changed the colors of the targets to orange and green. Control Phase 2 was presented at the end of the experiment to control for any effect of fatigue or globally reduced learning^41^ that might degrade adaptation. Finally, at the end of the experiment, a random trial was selected from one of the rewarded clamp blocks to determine participant bonuses. 15 out of 16 participants received the $10 bonus.

### Experiment 2 Protocol

Experiment 2 was aimed at both replicating the results of Exp. 1 and testing if the suppression of adaptation toward the punished target was object-referenced (due to punishment associated with the color of the target) or movement-referenced (tied to the particular reach directions associated with punishment in the later trials of the Reward Association Phase).

Experiment 2 began with the Reward Association Phase (**Fig. 1B**, *middle*). After learning the reward association, participants completed Test Phase 1 as described above. Prior to Test Phase 2, the Reward Association block was repeated, except with the target configurations now swapped (**Fig. 1B**). Thus, the final configuration of the rewarded and punished targets was maintained into Test Phase 2. This design choice was aimed at providing the most recent punishment history at the *new* location of the punished target, rather than at its original location, which was where the rewarded target would be placed in Test Phase 2. Importantly, the colors of the rewarded and punished target remained the same as the original Reward Association block. Participants then completed Test Phase 2 as in Experiment 1.

### Experiment 3 Protocol

Experiment 3 was aimed to test if adaptation was enhanced toward previously *rewarded* reach directions. Here, the values of the targets were modified such that the rewarded target always provided +10 points and was colored green. The other target, the “neutral” target, always provided +0 points and was shaded gray (**Fig. 1C**). Following the Reward Association Phase, participants were then instructed to always reach to the neutral target for the rest of the experiment. In Test Phase 1, participants went to the neutral target and the clamp blocks alternated the sign of the rotation such that adaptation proceeded either toward the rewarded target (TOWARD) or away from it (AWAY). In Test Phase 2, the target configuration swapped, such that the neutral target was now positioned where the rewarded target was previously, and participants were instructed to reach to the neutral target, in its new location.

### Data Analysis

The primary dependent measure was hand angle relative to the current goal target. In Experiment 1, hand angle was measured as the angular deviation between the reach direction and the goal target at peak movement velocity. Due to lower, less controlled sampling rates in the crowd-sourced experiments (Exps. 2 and 3), reach direction at endpoint was used. (We note that similar results were obtained in Experiment 1 when using endpoint hand angle).

Hand angles exceeding ±30° from the target were excluded. Trials where the participant received any of the previously mentioned warnings (i.e., too slow, too late, non-straight reach, reach to the wrong target) were also excluded. Participants for whom these criteria excluded 20% or more of their trials were excluded from further analysis. And participants who went to the wrong (i.e., non-cued) target for more than 10% of the rewarded washout trials with veridical feedback were also excluded, due to failure to comply with the instructions. No subjects were excluded from Experiment 1; 14 participants were excluded from Experiment 2, yielding a final sample of 33 participants; and 8 participants were excluded from Experiment 3, yielding a final sample of 36 participants. Overall, after excluding participants who failed to follow instructions, 4.9% of trials for the remaining participants were excluded in Exp. 1, 4.8% of trials in Exp. 2, and 4.8% of trials in Exp. 3.

To visualize adaptation learning curves, trials were binned in cycles of two trials, reflecting the number of potential target pair locations. All learning curves were “baseline-corrected” such that the average of the first two cycles was subtracted from all cycles in each block. This was especially important, as competing target values could simply induce a fixed movement bias instead of a modulation of adaptation. With this baseline correcting, learning within a block would thus reflect changes in hand angle accumulated over the block – actual adaptation – rather than reach angle drift that spanned across blocks or fixed biases.

Two methods were employed to quantify differences in adaptation between TOWARD versus AWAY blocks. First, we averaged the last two cycles within each block to get a measure of how much total (baseline-corrected) adaptation occurred within each block/phase for each participant. We then employed paired t-tests (or two-way ANOVAs) to quantify differences between adaptation in TOWARD versus AWAY blocks. We also performed exploratory nonparametric cluster permutation tests to visualize specific learning epochs where hand angles significantly differed between TOWARD and AWAY blocks^43,44^. To do this, we first performed a single paired one-tailed t-test on each cycle of the learning curves between TOWARD and AWAY blocks. If the initial paired t-test of the last two cycles showed significantly greater adaptation AWAY vs TOWARD in a given phase, this was used as the prediction in the one-tailed t-test of each cycle of that phase. If the initial comparison happened to be null, we assumed adaptation AWAY would be greater than TOWARD. A “cluster” was then defined as any string of adjacent significant cycles (p<0.05). For each cluster, we summed the t-statistic for each cycle to obtain a “t-sum statistic” for the cluster. We then computed a null distribution of t-sum statistics by randomly permuting the AWAY and TOWARD labels for each cycle 10,000 times. For each permutation, we identified significant clusters and computed the t-sum statistic for each permutation (if multiple clusters were identified, we used the one with the largest t-sum statistic). This generated a null distribution of t-sum statistics that would be generated by chance. If the t-sum statistic of a cluster in the original dataset was greater than the t-sum statistics of 95% of the permuted datasets, it was considered to be significant, thus controlling for the multiple comparisons used in this analysis.

In Experiment 1, we quantified repulsion effects as the difference between asymptotes of Test Phase 1 Away and Test Phase 1 Toward. We quantified suppression effects as the difference between asymptotes of Test Phase 1 Away and the average of the asymptotes for Test Phase 2.

### Model Fitting Procedure

Models were fit to each participant’s full reaching data in Test Phases 1 and 2 of all three experiments, including both clamp blocks and washout blocks, binned by cycles of 2 trials. In Experiment 1, model fitting also included the Control Phases, using each model’s baseline a and b terms to predict adaptation across all four Control Phase blocks. Parameter optimization was performed using Matlab’s *fmincon* function with 1000 maximum iterations. The objective was to minimize the root mean squared error (RMSE) of the hand angles predicted by the model from the subject’s true hand angles for all cycles. Models were compared using Bayesian Information Criterion (BIC). Parameter recovery for the Policy Gradient model and model confusability analysis for all 4 models was conducted and showed that the parameters of the Policy Gradient model were reliably recoverable and that the model was clearly dissociable from the alternative models (**Fig. S3**). Parameter comparison across experiments was conducted using Mann-Whitney U tests, using Matlab’s *ranksum* function. Reported |r| values are Pearson coefficient equivalents. Bayes Factors for null comparisons were computed using R’s *BayesFactor* package.

The Policy Gradient model was fit by first constructing the value landscape using the amplitude and loss aversion parameter to generate two Gaussians, one centered at the high-value target and one centered at the low-value target, each with a width of 20 degrees, and superimposing them on each other. This landscape reflected the learned reward “value” of all possible reach directions from −180 to +180 degrees (see Supplemental Note for the derivation of this value landscape from a policy gradient model). The 20 degree width was chosen because it matched the separation between the two targets. When allowed to be a free parameter, the vast majority of participants were fit by a Gaussian width within 5 degrees of 20 and the additional parameter was not justified by BIC. The resulting value landscape was then converted to a “learning rate modulation” value by adding 1 at all reach angles. On a given trial, the effective learning rate was found by selecting the modulation value for that reach angle and multiplying it by the baseline learning rate *b*_0_, with the resulting learning rate capped at a maximum of 1. Thus, learning rates varied on a trial-by-trial basis as the participants’ reach angles adapted to the error clamp. Finally, the value landscape decayed over time as the salience of the association likely diminished over time since the participant never reached to the punished target. This decay was modeled using a trial-by-trial decay factor *γ* < 1.

The Goal-Based Modulation model was fit by simply selecting a different learning rate for the TOWARD vs AWAY blocks. In Exps. 1 and 2, *b_A_* > *b_T_* (predicting repulsion from the low-valued target), while in Exp. 3 *b_A_* > *b_T_* (predicting attraction to the high-value target). This model predicted different learning rates based solely on the value of the “goals” or targets, rather than reach directions. Thus, it predicted that when adaptation brought the effector toward the low-value goal, adaptation would be suppressed, regardless of where the low-value goal was placed (i.e., it predicted a repelling effect in both Test Phases of Exp. 1). When adaptation brought the effector toward the high-value goal (in Exp. 3), the model predicted that adaptation would be enhanced (i.e., that adaptation would be biased toward the high-value goal).

The Bias Modulation model was fit by assuming a non-zero “hidden bias” at the start of each adaptation block. This bias was induced by the Reward Association phase (and Top-up in Exp. 2) and amounted to a flat clockwise or counterclockwise shift such that adaptation was skewed in the opposite direction from the lower valued target (negatively-valued target in Exps. 1 and 2; neutral target in Exp. 3). In TOWARD blocks, the bias meant that adaptation began closer to asymptote and thus only little further learning occurred before asymptote was reached. In AWAY blocks, learning began farther away from asymptote, and thus the observed learning appeared to be much greater. As in the Policy Gradient model, the bias decayed over time, to account for the apparently weaker effects observed in Test Phase 2.

### Data & Code Accessibility

Data files and analysis code for all experiments is available at https://osf.io/mvg97/overview?view_only=244c321be79a4d00bda87ed829ec268c

## Acknowledgements

This manuscript is the result of funding in whole or in part by the National Institutes of Health (NIH) [T32GM136577] and [R01NS132926]. It is subject to the NIH Public Access Policy. Through acceptance of this federal funding, NIH has been given a right to make this manuscript publicly available in PubMed Central upon the Official Date of Publication, as defined by NIH.

