## Supplemental Table S1, Supplement for "Instrumental-Motor Transfer: The Relative Value of Competing Movement Goals Modulates Implicit Motor Learning"

**Supplemental Materials**

**Supplemental Table S1**

| **Reward Association** | | | | | | |
| --- | --- | --- | --- | --- | --- | --- |
| Experiment | RT (ms, mean [SEM]) | | | MD (ms, mean [SEM]) | | |
|  | High Value | Low Value | p | High Value | Low Value | p |
| Exp. 1 | 574 (16) | 587 (20) | 0.18 | 92 (26) | 90 (24) | 0.5 |
| Exp. 2 | 456 (17) | 457 (19) | 0.94 | 330 (34) | 368 (48) | 0.3 |
| Exp. 2 (Top-up) | 436 (16) | 442 (17) | 0.53 | 103 (11) | 113 (12) | 0.43 |
| Exp. 3 | 458 (27) | 473 (21) | 0.47 | 495.3 (118) | 350 (64) | 0.24 |
| **Control Phases** | | | | | | |
|  | RT | | | MD | | |
|  | AWAY | TOWARD | p | AWAY | TOWARD | p |
| Exp. 1 CP 1 | 590 (17) | 582 (17) | 0.36 | 62 (8) | 50 (4) | 0.05 |
| Exp. 1 CP 2 | 567 (18) | 554 (14) | 0.11 | 58 (10) | 48 (3) | 0.22 |
| **Test Phases** | | | | | | |
|  | RT | | | MD | | |
|  | AWAY | TOWARD | p | AWAY | TOWARD | p |
| Exp. 1 TP 1 | 590 (15) | 582 (18) | 0.4 | 55 (6) | 54 (4) | 0.69 |
| Exp. 1 TP 2 | 582 (17) | 553 (14) | 0.003 | 53 (4) | 52 (3) | 0.62 |
| Exp. 2 TP 1 | 459 (19) | 449 (17) | 0.4 | 107 (12) | 105 (11) | 0.87 |
| Exp. 2 TP 2 | 456 (17) | 450 (18) | 0.43 | 80 (8) | 81 (7) | 0.93 |
| Exp. 3 TP 1 | 562 (76) | 464 (25) | 0.19 | 106 (12) | 104 (11) | 0.93 |
| Exp. 3 TP 2 | 465 (21) | 461 (25) | 0.71 | 80 (8) | 86 (10) | 0.55 |

**Table S1.** *Reaction times and movement durations across all experiments and phases.* All p-values from paired t-tests. SEM = standard error of the mean. RT = reaction time. MD = movement duration. High Value: RT/MD when reaching to the higher-valued target. Low Value: RT/MD when reaching to the lower-valued target.

**Supplemental Figure S1**

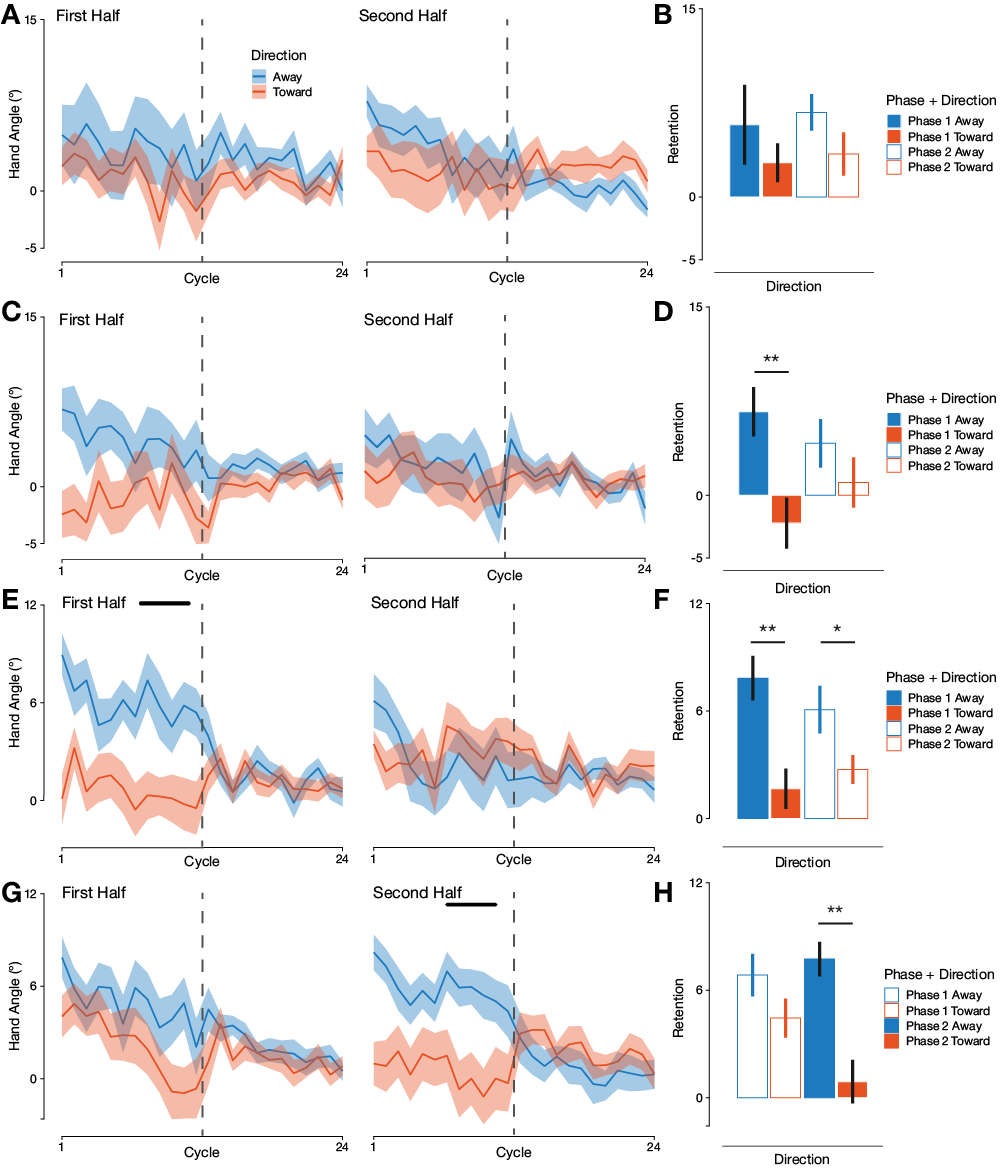

**Figure S1.** *Instrumental-Motor Transfer Retention in Washout Blocks.* **A)** Washout curves for Control Phase 1 (*Left*) and 2 (*Right*) in Exp. 1. **B)** Retention (baseline-corrected) in the Control Phases of Exp. 1. **C)** Washout curves for Test Phase 1 (*Left*) and 2 (*Right*) in Exp. 1. **D)** Retention in the Test Phases of Exp. 1. **E)** Washout curves for the Test Phases in Exp. 2. Bar indicates significant epochs (p<0.05) by cluster permutation test. **F)** Retention in the Test Phases of Exp. 2. **G)** Washout curves for the Test Phases in Exp. 3. Bar indicates significant epochs (p<0.05) by cluster permutation test. **H)** Retention in the Test Phases of Exp. 3. Dashed lines indicate the onset of veridical feedback during the washout. Error bars and shading reflect 1 S.E.M. * p<0.05 ** p<0.01.

Following each 48-trial block of adaptation to an error-clamp perturbation, participants completed 48-trial washout blocks consisting of 24 no-feedback trials and 24 veridical feedback trials. Here, we describe analyses of the results of the no-feedback washout trials across all phases of Experiments 1-3. The results largely parallel the results obtained from analyzing the adaptation trials.

*Method*

Washout retention was defined as the average of the first two cycles of the no-feedback washout, baseline-corrected by the first two cycles of the preceding clamp-adaptation block. Baseline-correction was done to ensure that washout retention was defined in such a way as to be comparable to our baseline-corrected adaptation metric used in analyses of the adaptation blocks. A two-way ANOVA was conducted on washout retention over Clamp Direction in the preceding block (TOWARD vs AWAY) and Phase (1 or 2). Further, paired t-tests were conducted between TOWARD and AWAY blocks in each phase. Finally, the results of these paired t-tests were used as the alternative hypothesis in exploratory cluster-permutation tests conducted over the whole no-feedback washout block, as was done in the clamped adaptation blocks. Cluster permutation tests were done to identify epochs within the washout block that significantly differed between TOWARD and AWAY conditions. Whenever the initial paired t-test was null, we assumed greater retention for TOWARD relative to AWAY as this was the hypothesized direction.

*Experiment 1 Washout Results*

In Test Phase 1, baseline-corrected washout retention was significantly reduced TOWARD the punished target relative to AWAY from it (8.9°, 95% CI [2.0, 15.7], t(15)=2.8, p=0.01, d_z_=0.69), replicating the results from the adaptation block. There were no significant differences between TOWARD and AWAY conditions in Test Phase 2 (t(15)-1.1, p=0.28) nor both Control Phases (CP1: t(15)=0.78, p=0.45; CP2: t(15)=1.3, p=0.22). Cluster permutation tests failed to reveal any significant epochs in both the Control Phases and Test Phases; however, one cluster was marginally significant in Test Phase 1 (t_sum_=7.5, p=0.067). Taken together, these results show that implicit adaptation was significantly attenuated TOWARD the punished target in Test Phase 1 and the effect persisted into the no-feedback washout block, further controlling for the use of any potential explicit strategies.

*Experiment 2 Results*

As in the adaptation blocks, baseline-corrected washout retention was significantly different between TOWARD and AWAY in both Test Phases, with significantly less retention TOWARD relative to AWAY (Test Phase 1: 6.2°, 95% CI [2.5, 9.8], t(32)=3.5, p=0.002, d_z_=0.60; Test Phase 2:, 3.3°, 95% CI [0.05, 6.6], t(32)=2.1, p=0.05, d_z_=0.36). The smaller effect in Test Phase 2 is consistent with the interpretation that the “top-up” Reward Association block (**Fig. 1B**) may have not fully abolished the initial learned association, leading to some interference that may have weakened the effect both in adaptation and washout. A two-way ANOVA on washout retention over Phase and Direction showed only a significant main effect of direction (F(1,128)=17.4, p=5.6x10^-5^), reflecting the finding that retention was significantly suppressed TOWARD relative to AWAY. Cluster permutation tests identified a significant cluster in Test Phase 1 (t_sum_=9.7, p=0.02), indicating that the washout curves significantly diverged during the no-feedback washout, with significantly more retention AWAY relative to TOWARD. Altogether, these results mirror and extend the findings from the adaptation blocks, suggesting that implicit adaptation is significantly attenuated in the vicinity of recently punished reach directions, and that this effect continued into the no-feedback washout blocks.

*Experiment 3 Results*

Washout retention was marginally less in the TOWARD condition relative to AWAY in Test Phase 1 (t(34)=2.1, p=0.05, d_z_=0.34). Mirroring the results from the adaptation blocks, retention was significantly suppressed in the TOWARD condition relative to the AWAY condition in Test Phase 2 (6.8°, 95% CI [3.4, 10.2], t(34)=4.0, p=0.0003, d_z_=0.68), and a cluster permutation test identified a significant cluster where washout curves significantly diverged in Test Phase 2 (t_sum_=11.1, p=0.03), indicating that implicit adaptation remained greater for AWAY relative to TOWARD into the no-feedback washout block. A two-way ANOVA on retention over Phase and Direction showed a marginally significant interaction (F(1,138)=3.9, p=0.05), indicating that the effect of clamp direction was significantly greater in Test Phase 2 than in Test Phase 1. These results parallel the results from the adaptation blocks and highlight that implicit adaptation was suppressed around relatively low-valued actions, even when those actions were not negatively-valenced in absolute terms. The fact that these effects persisted into the no-feedback washout further supports the conclusion that they reflect true implicit adaptation, rather than the use of explicit strategies.

**Supplemental Figure S2**

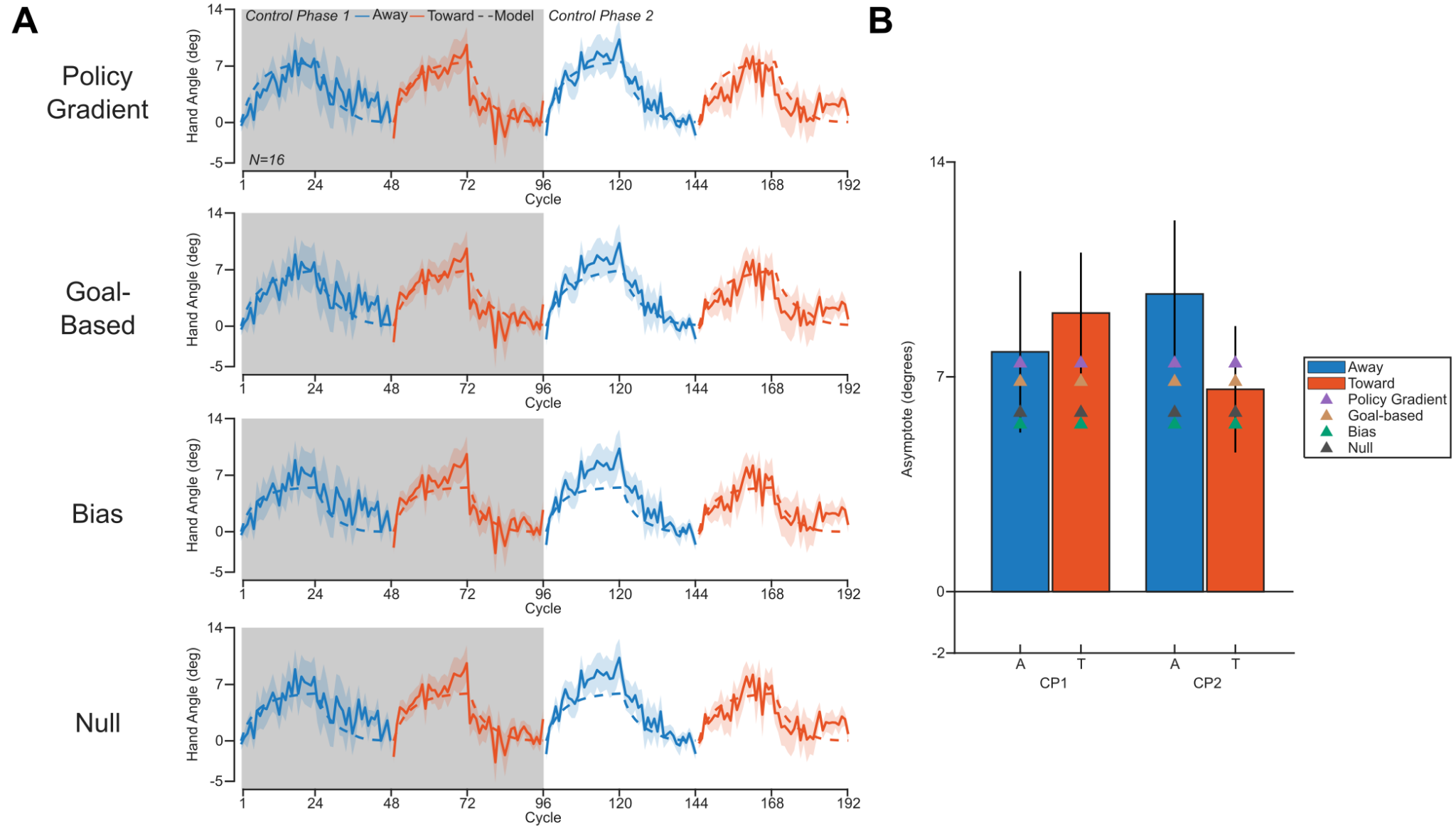

**Figure S2.** *Model fits to the Control Phases of Experiment 1.* **A)** Learning curves with associated model fits (dashed lines) for both Control Phases. Cycles averaged over 2 trials. Shaded regions reflect standard error of the mean. **B**) Asymptotes for both control phases with model-predicted asymptotes overlaid. Error bars reflect standard error of the mean.

**Supplemental Figure S3**

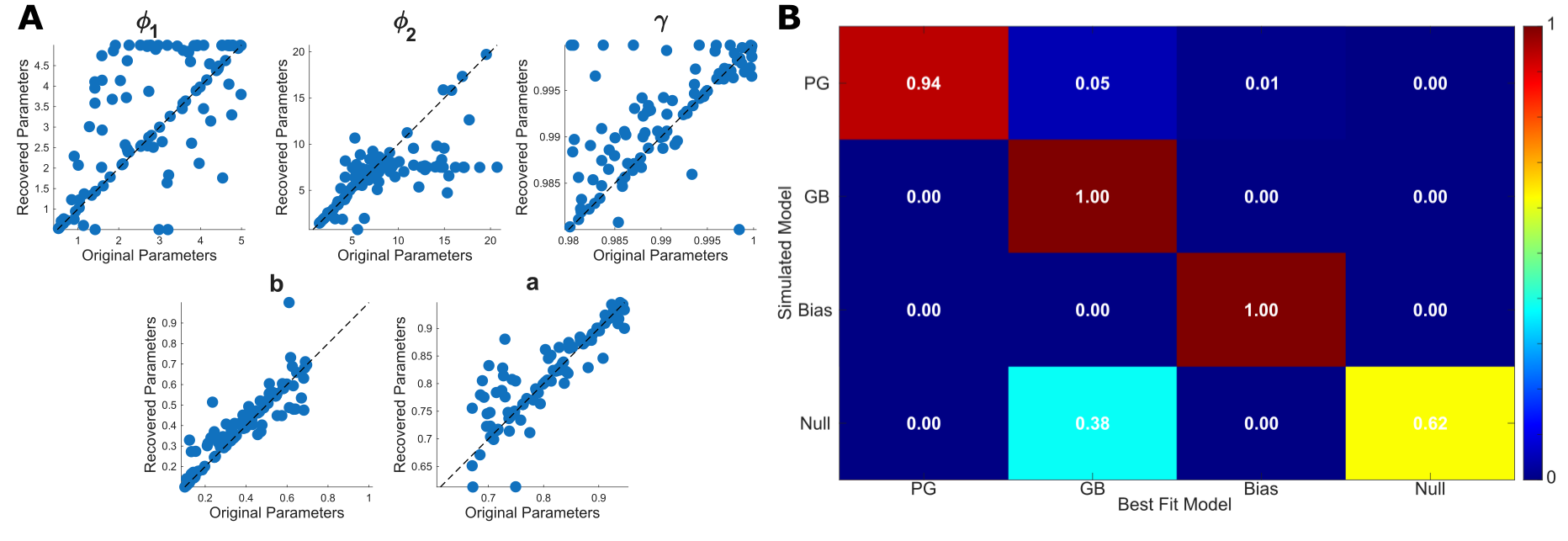

**Figure S3.** *Policy Gradient Parameter Recovery and Model Confusability.* **A**) Correlations between recovered and original parameters for all 5 parameters of the Policy Gradient model over 100 simulated agents. All correlations p<10^-12^. Average Spearman rho 0.75. Dashed line is the identity line. **B**) Model confusability matrix, showing the proportion of simulated agents best fit by their originating model.

*Parameter recovery and model confusability*

To ensure that our models were identifiable, and that the five parameters of the Policy Gradient model were recoverable, we conducted parameter recovery and model confusability analysis on simulated data from 100 agents for each of the four models. Agents were simulated over the range of parameters observed in Experiment 1, and we generated Control Phase and Test Phase learning curves for each agent as if they completed Experiment 1. We then fit each of the four models on the learning curves simulated for each of the 400 agents (100 for each model) and quantified how many agents were best fit by the model used to simulate their data vs another alternative model. The resulting confusability matrix is shown in Figure S3B. 100% of the Goal-Based Modulation and Bias Modulation agents were best fit by their original model. 94% of the Policy Gradient agents were best fit by their original model, with 5% better fit by the Goal-Based model and 1% better fit by the Bias model. 62% of the Null model agents were best fit by their original model, with 38% of agents better fit by the Goal-Based model.

We also conducted parameter recovery on the Policy Gradient model. We compared the parameters used to generate the simulated data to the parameters recovered when fitting the simulated data with the Policy Gradient model, to ensure that all parameters could be independently resolved. All recovered parameters significantly correlated with the original parameters (avg. ⍴=0.75, all p<10^-12^). Correlations for all 5 parameters are shown in Figure S3A.

**Supplemental Figure S4**

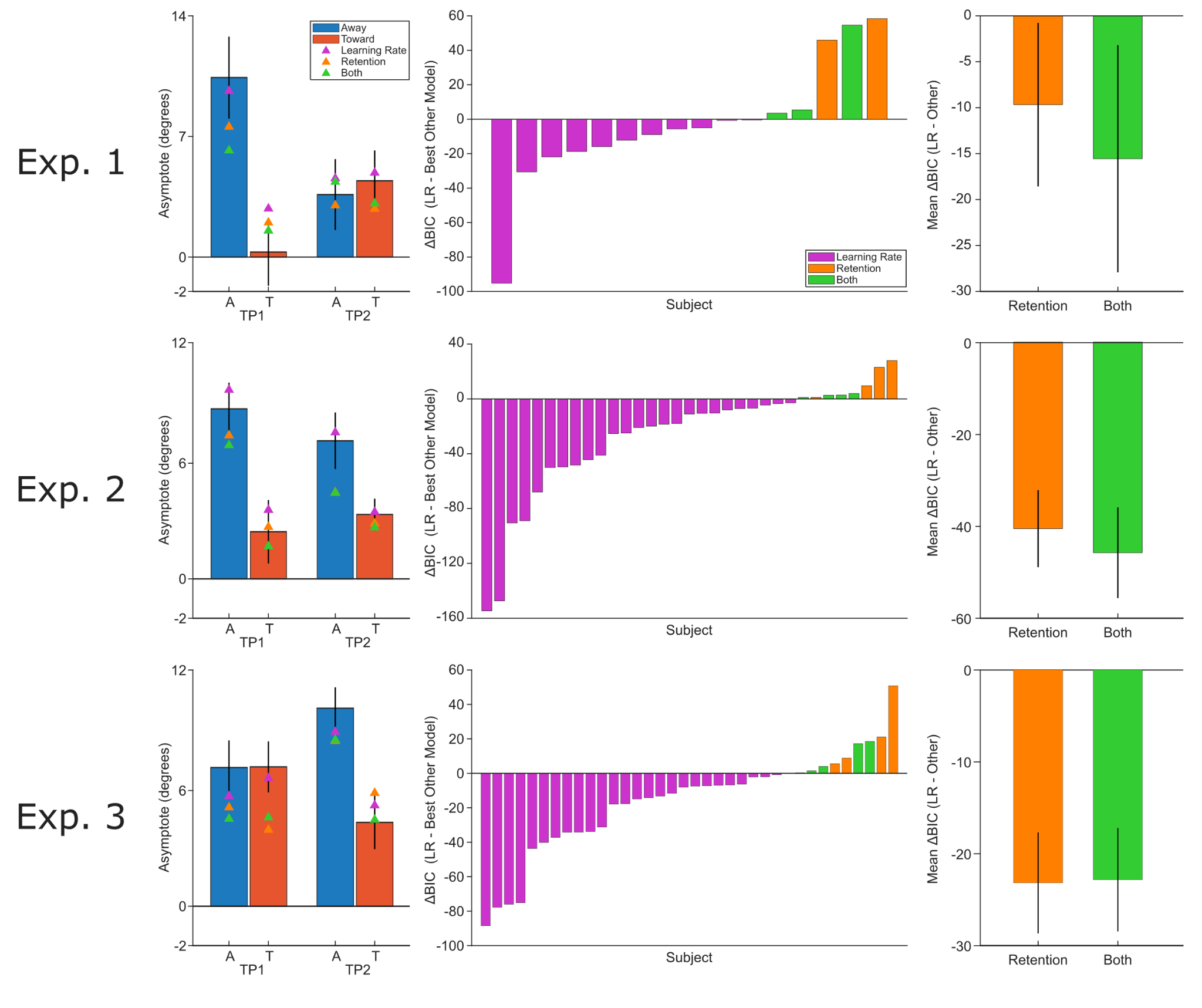

**Figure S4.** *Action reward values modulate learning rate, not other adaptation parameters. Left*: Asymptotes with model predicted asymptotes overlaid. Error bars reflect standard error of the mean. *Middle*: BIC difference between the Learning Rate model and the best alternative model (among Retention and Both). 11/16 participants were best fit by the Learning Rate model in Exp. 1 (*top*), 25/33 in Exp. 2 (*middle*), 25/36 in Exp. 3 (*bottom*). *Right*: Mean BIC difference between the Learning Rate model and the alternative variants. Error bars reflect standard error of the mean.

*Action-reward values modulate learning rate, not other adaptation parameters*

We fit participant data from all 3 experiments with variants of the policy gradient model where the learning rate, retention, or both parameters were modulated by the learned value landscape. Across all 3 experiments, the learning rate variant won by BIC (average ΔBIC vs retention, Exp. 1: -9.7, Exp. 2: -40.6, Exp. 3: -22.2; average ΔBIC vs both: Exp. 1: -15.6, Exp. 2: -45.9, Exp. 3: -19.5). In Experiment 1, 11/16 participants were best fit by the learning rate variant, with 2 best fit by the retention variant and 3 best fit by the variant that modulated both. 25/33 participants were best fit by the learning rate variant in Experiment 2, with 4 best fit by the retention variant and 4 best fit by the variant that modulated both. 25/36 participants were best fit by the learning rate variant in Experiment 3, with 3 best fit by the retention variant and 8 best fit by the variant that modulated both. Overall, 61/85 participants were best fit by the learning rate variant, or ~72%; thus, we conclude that action reward values modulate the learning rate of implicit adaptation, and not other adaptation parameters.

**Supplemental Figure S5**

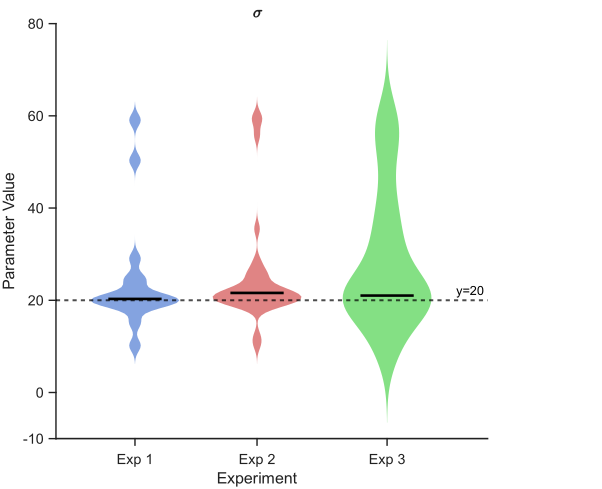

**Figure S5.** *Best-fit Gaussian widths in Experiments 1-3. Bar indicates median. Dashed line at 20 degrees.*

*Fitting the basis function width*

Adding a sixth free parameter to the policy gradient model to account for individual differences in the generalization width of the units was not justified by BIC across all 3 experiments (average ΔBIC, Exp. 1: 0.45, Exp. 2: 1.29, Exp. 3: 3.4). 12/16 participants in Experiment 1, 23/33 in Experiment 2, and 24/36 in Experiment 3, had a best fitting width that was within 5 degrees of 20. The best-fit width parameter averaged 24 ± 13 in Exp. 1, 25 ± 12 in Exp. 2, and 24 ± 12 in Exp. 3. Since ~70% of the participants had a best fit width within 5 degrees of 20, and the average best-fit width was within 5 degrees of 20, we conclude that setting the width to 20 degrees, equal to the separation between the two targets, was reasonable.

**Supplemental Note**

*Theoretical basis of the policy gradient model*

To model adaptation across all three experiments, we adopted a policy gradient model which learned the value of all actions (reach directions) and converted this to an implicit adaptation learning rate for a given reach direction. This architecture required two distinct modules: a reinforcement learning (RL) module, modeled after an actor-critic model of the basal ganglia (Greenstreet et al. 2026), and a motor adaptation module, employing the popular state-space model of implicit adaptation (Throughman & Shadmehr, 2000), thought to be implemented by the cerebellum.

The RL module consists of two agents: the “critic”, a discrete Q-learning agent that learns the values of each of the two competing movement goals, and the “actor”, a policy gradient agent that learns the values of all possible actions (i.e., reach directions). During the reward association phase, the critic samples the outcomes of hitting either of the two targets and learns the value associated with each of them, updating the value of the goal by a proportion of the reward-prediction error (RPE) experienced on that trial.

$\delta(t)=r(t)-Q_{i}(t) if r(t)>0$	(1)

$=\lambda\cdot r(t)-Q_{i}(t) if r(t)<0$

$Q_{i}(t+1)=Q(t)+\alpha\cdot\delta(t)$	(2)

Here, $Q_{i}$ reflect the value of each of the two targets, $\delta$ is the RPE, $r$ is the reward received, $\lambda$ is the loss aversion parameter that scales punishments by a scalar factor, and $\alpha$ is the critic’s learning rate.

The learned values for each of the targets are then transformed into relative values before being transmitted to the actor to guide the actor’s learning of the value of all possible reach directions.

$V=Q_{goal}-Q_{nongoal}$	(3)

where V is the relative value of the target the agent hits compared to the other non-goal target. On trials where the agent misses both targets, V would be 0 as neither target was hit and it is difficult to infer which target was the true “goal”.

The “actor” is modeled as a policy gradient agent. Policy gradient models are a subset of direct policy learning models that differ from discrete RL models in that agents directly learn a continuous valued “policy” across all actions, rather than discrete values for a finite number of actions. To do so, we first parameterize the action space of reach angles, creating basis units $\theta_{i}$that are tuned such that they are most active to a preferred reach direction but have a broad tuning curve for adjacent reach angles.

$\theta_{i}(x)\sim N(\theta_{preferred}, \sigma^{2})=e^{{(x-\theta_{pref})}^{2}/\sigma^{2}}$	(4)

The exact number of units is arbitrary, as long as there are enough units to tile the range of possible reach directions since the units are highly overlapping due to their broad tunings. The tuning width of these basis units $\sigma^{2}$was chosen to be 20 degrees as this approximates a typical width for such generalization functions and was the motivation behind separating the targets by 20 degrees.

We then instantiate a weight vector $\omega_{i}$, representing the weight applied to each of the basis units, or, in our case, the value of each particular reach direction. Initially, each $\omega_{i}=0$. However, on each trial, the critic provides the actor with the relative value information specific to that trial to update the value of all the reach directions proportional to the activation of each of the basis units for the reach direction that was actually executed:

$\omega(t+1)=\omega(t)+\eta\cdot V\cdot\theta(x)$	(5)

where $\eta$ is the actor’s learning rate, $V$ is the relative value from Eq. 3, and $\theta(x)$ is the vector representing the activation of each of the basis units evaluated at $x$, the true reach angle on that trial. Thus each unit’s weight is increased by an amount proportional to its activation if that reach angle was rewarded and decreased by an amount proportional to its activation if that reach was punished.

Typically, when employing a policy gradient model, the policy is constructed from the convolution of the weights with the basis units. However, in our task, participants are explicitly instructed to go to one of the two targets, so their choice policy is effectively ignored.

Over the course of the Reward Association phase, the RL module learns a “value landscape” $\omega$, representing the value of each reach direction (i.e. each $\omega_{i}$ corresponds to a basis function with preferred reach direction $\theta_{i}$). Since we are not interested in the temporal evolution of this landscape during the Reward Association phase, and we are only interested in the final learned value landscape, we model the landscape as two Gaussians centered at the good and bad target, with widths corresponding the basis unit widths and amplitudes $\varphi_{1}>0$ and $\varphi_{2}<0$, corresponding to the good and bad targets, respectively. We define a new “loss aversion” parameter as the ratio $\lambda^{*}=\left| \varphi_{2}/\varphi_{1} \right| > 1$. This simplification is equivalent to the full policy gradient model with actor-critic trained on the exact Reward Association phase used in the task (see code in Data & Code Accessibility), thus in our modeling we used this simplified method to construct the value landscape.

In Test Phases 1 and 2, this value landscape is then converted to an adaptation “learning rate” (or error sensitivity) by multiplying by a “baseline learning rate” $b_{0}$ by the weight plus 1 and capping the resulting learning rate at 1, as learning rates cannot exceed 1 by convention.

$b(x)=min(b_{0}\cdot(\omega(x)+1), 1)$	(6)

Thus, learning rates are no longer static, but vary as a function of the reach angle during adaptation. The +1 in Eq. 6 is to account for the fact that all weights start at zero. If no learning occurred, all learning rates would be the baseline learning rate $b_{0}$. Reach angles are then adapted by the classic state space equation:

$x(t+1)=b\cdot e(t) + a\cdot x(t)$	(7)

where $e(t)$ is the motor error experienced on the trial (e.g., the clamp magnitude during clamped trials) and $b$ is the retention term, reflecting how quickly adaptation washes out in the absence of motor errors. $b$ is derived from Eq. 6. We tested models where the value landscape $\omega$ modulated retention $a$ or both retention and learning rate parameters separately. The results of this modeling are included above and highlight that the value landscape appears to modulate the learning rate and not other adaptation parameters (see “*Action-reward values modulate learning rate*” above; Fig S4).

During Test Phases 1 and 2, the learned weights are subject to decay as the participants never reach to the punished target and the salience of the association diminishes over time. For Exps. 1 and 3 this is modeled as an exponential decay:

$\omega(t+1)=\omega(t)\cdot\gamma$	(8)

Where $\gamma$<1. This decay term explains the apparent diminishing of the effect in TP2 relative to TP1 and reflects the assumption that the weights diminish to zero over a long enough timescale (i.e., that adaptation eventually returns to normal).

For Experiment 2, where there is an intervening “top-up” Reward Association phase (reversal), the learned value landscape must be updated. To do so, we swap the exponential decay parameter for an “interference” $\kappa\in[0,1]$ parameter. $\kappa$ multiplies the original value landscape to reflect degradation over the course of TP1 before adding the new learned value landscape (with the locations of the good and bad target swapped) to obtain the final resultant value landscape. This reflects the assumption that the learned value landscape decays subtly over time, but is not completely abolished before the “top-up” Reward Association Phase. This simplification closely replicates a full trial-by-trial simulation of the policy gradient model (see code in Data & Code Accessibility). Kappa can be converted to a trial-by-trial decay parameter $\gamma$ by exponentiating to the 1/96th power. This trial-by-trial decay was used to decay the weights across both Test Phases.

Thus, our model has 5 tunable parameters: $\varphi_{1}$ and $\varphi_{2}$ reflect the amplitude of the value landscape at the good and bad target respectively; $b_{0}$ and $a$ reflect the baseline learning rate and retention parameters for the adaptation module; and $\gamma$ (Exps. 1 and 3) or $\kappa$ (Exp. 2) reflect the decay of the value landscape over time. Allowing the basis unit width $\sigma^{2}$ to be a tunable parameter did not significantly change our fits and was not justified by BIC analysis, and resulted in the majority of participants being fit by a width of 20˚ ± 5˚ for all 3 experiments (see “*Fitting the basis function width*” above; Fig. S5).
